# Projection-specific Routing of Odor Information in the Olfactory Cortex

**DOI:** 10.64898/2025.12.12.694045

**Authors:** Simon Daste, Tuan H. Pham, Max Seppo, Alexandre André, Shyam Srinivasan, Jingyun Xiao, Andrea Sattin, Chiara Nardin, Tommaso Fellin, Kevin M. Franks, Eva Dyer, Alexander Fleischmann

## Abstract

Sensory processing in the mammalian cortex relies on extensive feedforward and feedback connections, yet how information is routed along these pathways remains poorly understood. Here, we examined the functional properties of feedback and feedforward neurons in the mouse olfactory (piriform) cortex. We selectively labeled neurons projecting to the olfactory bulb (OB, feedback) or medial prefrontal cortex (mPFC, feedforward) and recorded their activity during passive odor exposure and learning of an odor discrimination task. We found that odor identity and reward associations were encoded by OB-projecting ensembles early during odor exposure, whereas mPFC-projecting neurons encoded this information later, aligned with behavioral responses. Moreover, mPFC-projecting neurons maintained a stable representation of valence across days, while OB-projecting neurons exhibited pronounced plasticity. Together, these findings reveal that odor information is selectively routed through feedforward and feedback pathways and suggest that the functional properties of piriform neurons mirror the computational demands of their downstream targets.

## Introduction

Sensory systems transform physical features of the environment into actionable percepts. Sensory processing occurs along hierarchically structured neural pathways that carry stimulus-evoked neural activity from the periphery to higher-order brain regions. In mammals, feedforward pathways transform basic stimulus features, for example edges in vision, spatial location in touch, or frequency in audition, along the cortical hierarchy into more complex representations of sensory objects and scenes (1–6). Along the same pathways, feedback projections dynamically reshape sensory representations, including by regulating gain and incorporating contextual and attentional information (7–13). Hence, many brain areas exhibit contextually modulated tuning to increasingly complex features across the hierarchy of sensorimotor transformation. In vision, for example, neurons in the primary visual cortex of primates selectively respond to basic stimulus features such as edge orientation, while neurons in the inferotemporal (IT) cortex represent complex features including visual objects and scenes (14–16). At the same time, feedback signals to IT support accurate object recognition (17) and visual episodic memory formation (18).

This bidirectional, hierarchical model of information processing implies that within a given brain region, feedforward and feedback projection neurons transmit distinct stimulus features, often conceptualized as error signals and top-down predictions in predictive coding frameworks (19,20). However, the functional specializations of feedforward and feedback projection neurons along hierarchical neural pathways and how they are shaped by learning and experience remain poorly understood.

To address this question, we leveraged the compact organization of the mammalian olfactory system. Unlike other sensory systems in which early sensory processing involves multiple stations including obligatory thalamic relays, inputs from olfactory receptor neurons in the periphery can reach the olfactory (piriform, PCx) cortex directly via the olfactory bulb (OB) (21–24). PCx sends extensive feedback projections back to the OB, while feedforward pathways project directly to higher-order associative areas such as the medial prefrontal (mPFC) and lateral entorhinal cortex (25–32). Furthermore, learning and experience modulate the functional properties of piriform neurons, and PCx has been shown to be critical for olfactory learning and memory (22,33–37). Thus, olfaction provides an attractive model to explore how sensory information is processed along hierarchically structured neural pathways and how this processing is shaped by learning and experience.

Here, we asked whether PCx neurons projecting to the OB (feedback) and mPFC (feedforward) exhibit distinct functional properties. We used two-photon calcium imaging in awake mice to record odor-evoked responses from each projection neuron type, during both passive odor exposure and while mice learned to perform an odor discrimination task. We found that OB-projecting neurons represented odor identity, concentration, and reward associations rapidly upon odor exposure, suggesting a role in fast, feedback-driven refinement of sensory input. In contrast, mPFC-projecting neurons exhibited delayed encoding of odor identity and reward associations, coinciding with the animal’s behavioral response. Together, these results reveal a functional segregation of odor information routing between feedforward and feedback neural pathways.

## Results

### Calcium imaging of piriform neurons with distinct projection targets

To investigate odor coding properties of OB- and mPFC-projecting piriform cortex (PCx) neurons, we injected an Adeno-Associated Virus (AAV) driving pan-neuronal expression of jGCaMP7f (38) into the anterior PCx (aPCx). Additionally, we injected a retrogradely transported AAV (AAV-retro; Tervo et al. 2016) expressing Cre recombinase into either the granule cell layer of the OB or the infralimbic/prelimbic regions of the mPFC of a Cre-dependent tdTomato reporter mouse line (Ai14; (40); **Figure 1A, Figure S1** and **Methods**). We then implanted an aberration-corrected gradient index (GRIN) lens-based endoscope (41) above the aPCx to enable chronic in vivo two-photon imaging in awake, head-fixed mice (**Figure 1B** and **C, Figure S1**). During two-photon imaging, we reliably detected GCaMP and tdTomato co-expression, defining OB-p and mPFC-p neurons, while untagged neurons with unidentified projection targets were marked by GCaMP expression alone (**Figure S2**).

**Figure 1.**
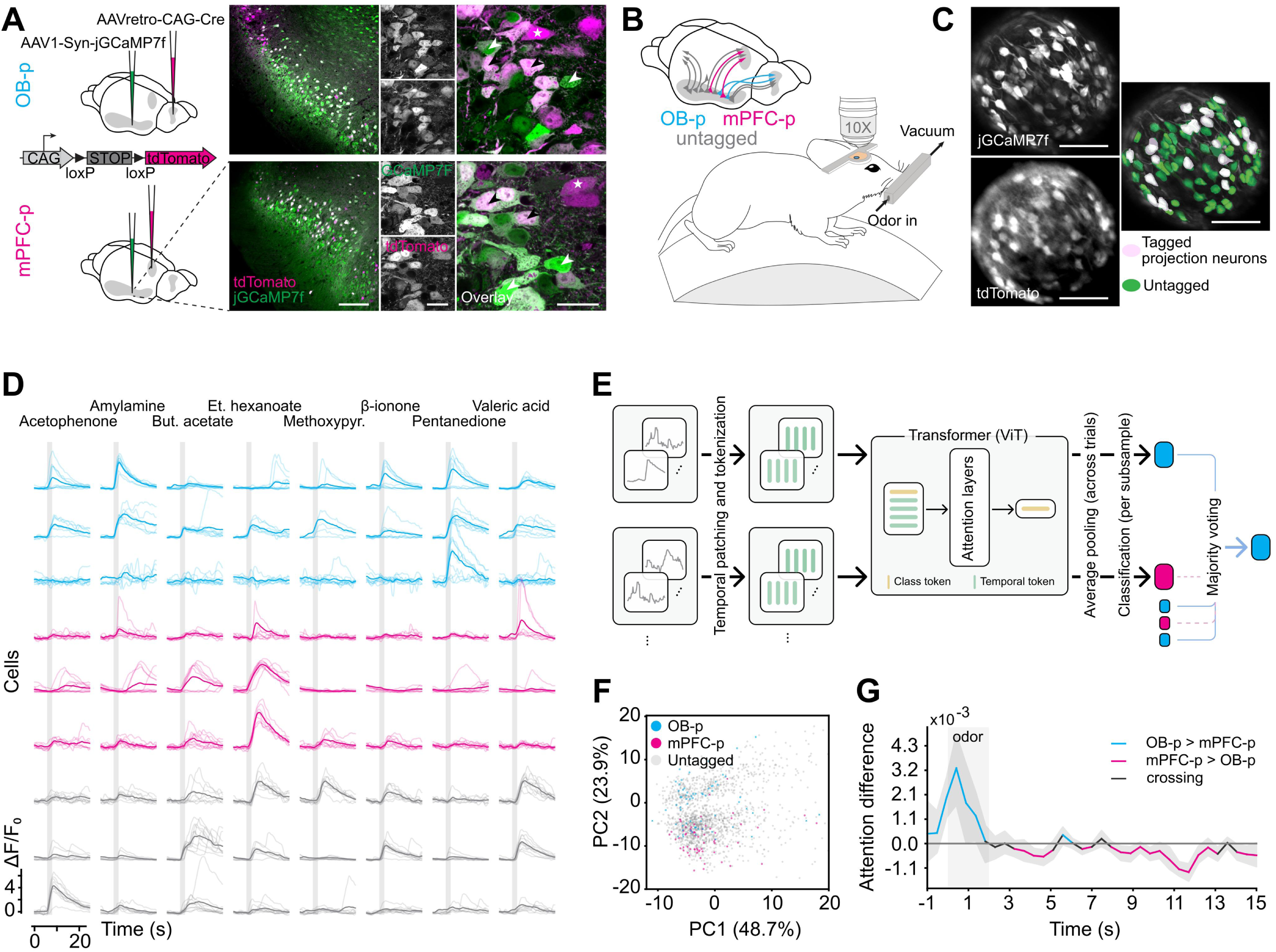
Calcium imaging reveals projection-specific activity patterns and temporal signatures that predict OB-p versus mPFC-p identity. (**A**) Schematic of viral labeling. AAV1-Syn-jGCaMP7f was injected into the PCx to achieve pan-neuronal expression of jGCaMP7f, and AAVretro-CAG-Cre was injected into either the olfactory bulb (OB; top) or medial prefrontal cortex (mPFC; bottom) of Ai14 mice to selectively express tdTomato in OB- or mPFC-projecting neurons. *Right:* Representative coronal section from the PCx showing widespread jGCaMP7f expression (green) and sparse tdTomato labeling (red) of the same section. White arrowheads: jGCaMP7f-only neurons. Black arrowheads: double-labeled neurons. White stars: tdTomato-only neurons. Scale bar (overview): 200 μm; scale bar (zoom-in): 50 μm. (**B**) Top: Schematic showing the three populations investigated: PCx (untagged), OB-p, and mPFC-p. Bottom: Schematic of the experimental setup. (**C**) Example images show the maximum projection of jGCaMP7f (top) and tdTomato (bottom) fluorescence signals through the GRIN lens, segmented ROIs, and their overlay. Double-positive ROIs (light purple) indicate labeled projection neurons. (**D**) Representative calcium traces. Example ΔF/F₀ time series from OB-projecting (cyan), mPFC-projecting (magenta), and untagged (gray) neurons. (**E**) Simplified schematic of the transformer-based classifier for predicting neuron projection targets from single-cell calcium dynamics. Calcium traces from individual neurons were sampled across trials to form subsamples. Within each subsample, trials were divided into non-overlapping temporal patches and tokenized for input to a transformer (ViT-style). Trial-level latents were averaged, pooled, and passed through a classification head to predict projection identity. Final neuron-level classification was obtained by majority voting across subsamples. See **Methods** and **Figure S3A** for more information and detailed schematics. (**F**) Principal component analysis (PCA) of latent representations extracted by the transformer model trained on the combined passive identity and Go/NoGo datasets (see **Methods**) and evaluated on the former (see **Figure S3F** for PCA with the latter). Each point corresponds to the latent embedding of a single cell (see also **Figure S3E** for differences between representations of subsamples and single cells), colored by projection targets (cyan: OB-p, magenta: mPFC-p, grey: untagged) and classification outcomes (circle: correct, cross: incorrect; for known projection targets). (**G**) Attention-rollout analysis of the transformer model trained on the combined passive identity and Go/NoGo datasets then evaluated on the former (see also **Figure S3D** for evaluation on the Go/NoGo dataset). The plot shows the temporal difference (averaged across 10 seeds, shading: 95% CI) in attention weights between correctly classified subsamples corresponding to their true neuron’s projection labels (OB-p vs mPFC-p). For each subsample, attention values were averaged across trials. Positive values (cyan) indicate time windows where attention was stronger for OB-p neurons, whereas negative values (magenta) correspond to higher attention for mPFC-p neurons. Black lines denote crossing epochs. Vertical gray area: odor presentation.

We generated two complementary datasets using this experimental approach. In the first (passive) dataset, head-fixed mice were either exposed to 8 monomolecular odorants at a single concentration (10 mice, totaling 81 OB-p, 84 mPFC-p, and 1978 untagged aPCx neurons, **Table 2**), or to three odorants at three concentrations each (10 mice, totaling 87 OB-p, 102 mPFC-p, and 2232 untagged cells, **Table 3**). In the second Go/NoGo (GNG) dataset, 7 mice (4 OB-p–tagged, 3 mPFC-p–tagged) performed a 4-odor discrimination task in which two odorants previously associated with water reward served as CS+ and two novel unrewarded odorants served as CS– (**Table 4**). Per session (mean ± SD), we recorded 95 ± 2 OB-p, 87 ± 8 mPFC-p, and 1690 ± 35 untagged neurons.

### Temporal structure of activity predicts OB-p vs mPFC-p identity

We first asked whether the calcium dynamics we recorded could distinguish OB-p from mPFC-p neurons. We trained a multilayer perceptron (MLP) on ΔF/F traces (**Figure 1D**) pooled across the odor-identity (passive) and Go/NoGo datasets (see **Methods**). The MLP achieved F1 scores of 0.688 (OB-p) and 0.557 (mPFC-p), providing initial evidence that the two projection targets could be differentiated based on their neural activity patterns alone. Next, we designed a transformer-based model (**Figure 1E, Figure S3A**) that leverages temporal attention, which improved test performance to 0.759 (OB-p) and 0.744 (mPFC-p) (**Figure S3B**). As a more stringent test of performance, we held out all projection-tagged neurons from one OB-p and one mPFC-p animal for testing. Performance remained high (F1: 0.754 OB-p; 0.623 mPFC-p), indicating high generalization capacity of the transformer to decode projection targets based on neural activity (**Figure S3B**).

To investigate the aspects of neural activity that the model uses to classify projection targets, we first extracted the latent embeddings from the transformer’s penultimate layer, which separated OB-p and mPFC-p well with minimal clustering by animal or dataset (**Figure 1F** and **Figure S3E-G**). Next, we examined the temporal attention map, using an attention rollout method (42) on correctly classified cells, which indicated that informative epochs cluster early after odor onset (around 0-3 s) for OB-p neurons, but later in the trial for mPFC-p neurons (see **Figure 1G** for the identity dataset and **Figure S3D** for the Go/NoGo datasets).

These results suggest that the temporal structure of neural activity can distinguish neural sub-populations defined by their projection targets.

### OB-p and mPFC-p neurons exhibit broadly similar odor response properties, but differ in dynamics

Previous studies have shown that different odors activate sparse, partially overlapping ensembles of PCx neurons (43–47). Thus, we first examined basic odor responses OB-p and mPFC-p neurons, in mice passively exposed to 8 different odorants (**Figure 2A-E**).

**Figure 2.**
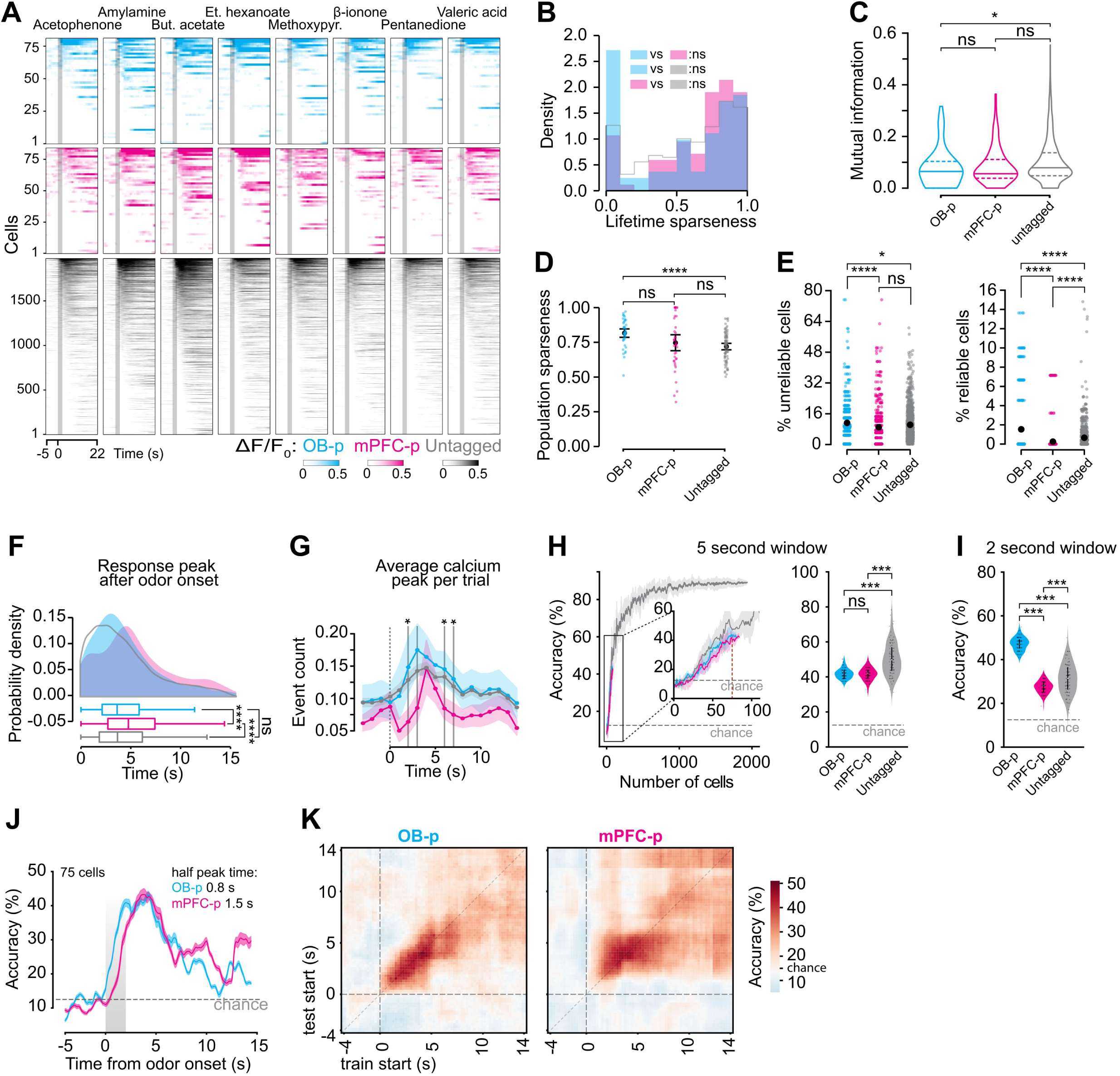
Differential odor response timing and selectivity distinguish OB- and mPFC-projecting piriform neurons. (**A**) Heatmap of trial-averaged odor responses. Each row corresponds to a neuron, sorted independently within OB-p, mPFC-p and untagged groups (y-axes) for each odor. Vertical red dashed lines indicate odor onsets. (**B**) Lifetime sparseness distributions for OB-p (n = 81), mPFC-p (n = 84) and untagged (n = 1978) neurons. (**C**) Mutual information between binarized activity and odorant identity in OB-p (n = 81), mPFC-p (n = 84) and untagged (n = 1978 neurons) groups. (**D**) Population sparseness per trial for OB-p, mPFC-p and untagged groups. Points represent single trials (n = 64). (**E**) Percentage of responsive PCx cells that are unreliable (left) and reliable (right) per trial in OB-p (n = 81), mPFC-p (n = 84) and untagged (n = 1978 neurons) groups. (**F**) Distribution of latencies to first calcium activity peaks after odor onset for OB-p (n = 24), mPFC-p (n = 27) and untagged (n = 424) neurons. (**G**) Average number of calcium activity peaks in a trial, aligned to odor onset for OB-p (n = 24), mPFC-p (n = 27) and untagged (n = 424) neurons. (**H**) Odor identity classification using a linear SVM, trained and tested on response vectors from a 5 s window after odor onset. Left: Classification accuracy using pseudo-populations of increasing size. Single-trial responses to 8 odors were decoded using a linear SVM classifier. Mean accuracy (± shaded error region) is shown across 100 random subsampling iterations of cells, with each subsampling evaluated over 10 independent train/test repeats. Dashed line indicates the chance level (12.5%). Inset: zoomed in view on 100 neurons. Right: Classification accuracies for 200 train/test repeats with matched population sizes (75 cells) across pseudo-population types. (**I**) Same as (**H**) except classifiers were trained and tested using a 2 s window after odor onset. (**J**) Odor identity classification accuracy of a linear SVM, trained and tested on response vectors from 250 ms sliding windows, using pseudo-populations of 75 cells (100 repeats). Dashed line indicates chance level (12.5%). Shading around lines: 95% CI. (**K**) Odor identity classification accuracy of a linear SVM, trained (x-axis) and tested (y-axis) on response vectors from non-overlapping 250 ms windows, using pseudo-populations of 75 cells, with 100 train/test repeats. For all figures, unless otherwise noted, the level of statistical significance is denoted as: ns: p > 0.05; *: 0.01 < p <= 0.05; **: 1 × 10⁻^3^< p <= 1 × 10⁻^2^; ***:1 × 10⁻^4^< p <= 1 × 10⁻^3^; ****: p <= 1 × 10⁻^4^.

Odor responses in PCx were generally sparse on the single cell level, with no significant differences in lifetime sparseness between groups (**Figure 2B**). Mutual information did not differ significantly between projection sub-populations, but was lower for OB-p neurons than for untagged neurons (p = 0.012) (**Figure 2C**). At the ensemble level, population sparseness differed only between OB-p ensembles and the untagged populations (**Figure 2D**, p = 1.26 × 10⁻^5^). The two sub-populations differed significantly in the proportion of cells that responded to odorants, with OB-p neurons responding in a significantly higher proportion than mPFC-p neurons (two-sided Mann–Whitney U test with Bonferroni corrected p-values, p = 5.79 × 10⁻^7^). We also quantified trial-to-trial response reliability, classifying neurons as reliable if they responded on >50% of odor presentations, and found that OB-p populations contained higher fractions of both unreliable and reliable cells (10.97% and 1.54%, respectively) than mPFC-p populations (**Figure 2E**, 8.71% and 0.25%; two-sided Mann–Whitney U test with Bonferroni corrected p-values: unreliable p = 3.90 × 10⁻^4^, reliable p = 8.20 × 10⁻^14^). Overall, these data suggest that while both OB-p and mPFC-p neurons are canonically odor responsive, OB-p neurons respond more strongly and more selectively than mPFC-p neurons.

To probe response dynamics, in a subset of experiments (5 mice, totaling 24 OB-p, 27 mPFC-p, and 424 untagged cells), we examined the temporal structure of odor responses by performing high-speed (30 Hz) imaging of odor presentations aligned to the inhalation phase of the first sniff. We found that, on average, OB-p neurons exhibited shorter latencies to reach the first peak of calcium activity following odor onset compared to mPFC-p neurons (**Figure 2F**; two-sided Mann-Whitney U test with Bonferroni corrected p-values: OB-p vs mPFC-p = 1.18 × 10⁻^17^, OB-p vs untagged = 1, mPFC-p vs untagged = 5.05 × 10⁻^25^). OB-p neurons also had more peaks during earlier time points than mPFC-p neurons (**Figure 2G**, and **Figure S4A-C**).

Next, we compared odor encoding dynamics. Odor identity could accurately be decoded from OB-p and mPFC-p pseudo-populations. Using linear SVM classifiers on size-matched pseudo-populations trained on activity within a 5-second window, decoding accuracy increased with population size for all groups, and both OB-p and mPFC-p pseudo-populations reached similar accuracy levels, significantly above chance but slightly lower than the untagged population (**Figure 2H**; two-sided Mann-Whitney U test, p = 0.398).

SVM classification using a sliding window for odor responses revealed that decoding accuracy for OB-p neurons peaked earlier than for mPFC-p neurons (**Figure 2J**). Consistent with this finding, when we compared odor decoding in pseudo-populations of increasing size in an early window (0-2 seconds), we found that OB-p pseudo-populations outperformed mPFC-p pseudo-populations (**Figure 2I**, two-sided Mann-Whitney U test with p = 2.15 × 10⁻^30^). Interestingly, decoding analysis with a sliding window also shows that mPFC-p pseudo-populations maintained low but above-chance identity decoding that exceeded OB-p decoding later in the trial (**Figure 2K**).

Together, these results suggest substantial overlap in the odor response characteristics of OB-p feedback and mPFC-p feedforward PCx neurons, but divergence in their response dynamics.

### Odor concentration is robustly encoded in OB-p but not mPFC-p neurons

Given the important role of the OB in gain control (48–50) we next asked whether OB-p feedback neurons preferentially encode odor concentration compared with mPFC-p feedforward neurons. To address this, we recorded responses to an odor panel consisting of three odorants, each presented at three concentrations spanning two orders of magnitude (10 mice, totaling 87 OB-p, 102 mPFC-p, and 2,232 untagged cells; **Table 3**; see also **Figures 3** and **S5**). Example traces illustrate that some OB-p neurons showed graded changes in response amplitude across concentrations for a given odor, whereas mPFC-p neurons often exhibited modest or saturating responses over the same range (**Figure 3A**).

**Figure 3.**
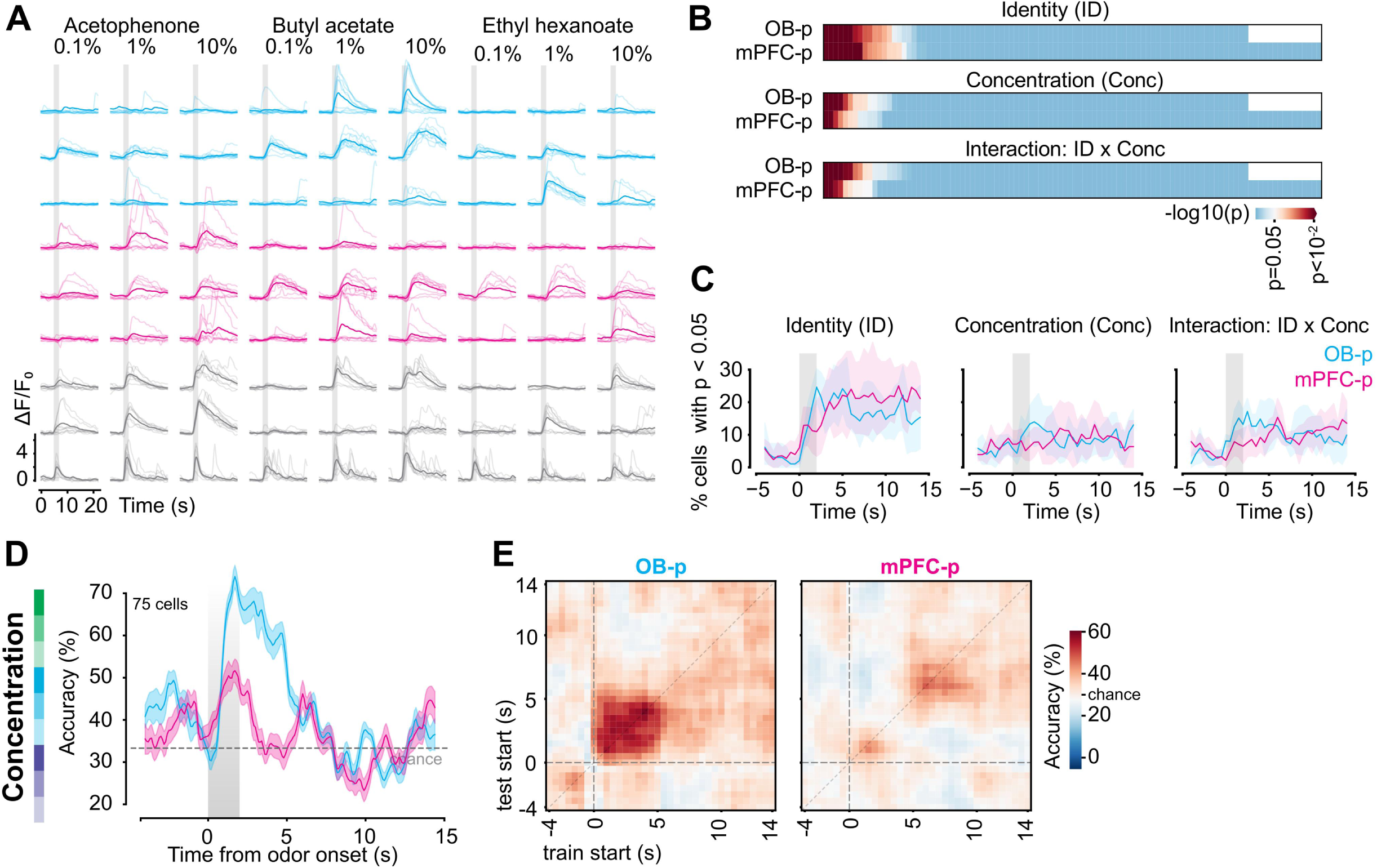
Odor concentration is robustly encoded in OB-p but not mPFC-p neurons. **(A**) Example calcium responses from individual OB-p (cyan), mPFC-p (magenta), and untagged (gray) neurons to the three odorants (acetophenone, butyl acetate, and ethyl hexanoate) presented at three concentrations spanning two orders of magnitude (0.1%, 1%, and 10%). Darker traces show trial-averaged ΔF/F₀. (**B**) Single-neuron two-way ANOVA (odor identity × concentration) applied to mean fluorescence during the first 5 s of odor presentation. Heatmaps show –log₁₀(p) values for significant effects of identity (ID), concentration (Conc), or the identity × concentration interaction. (**C**) Two-way ANOVA quantifying the proportion of neurons significantly modulated by identity, concentration, or their interaction across time. For each projection type, the percentage of neurons with p < 0.05 is shown in sliding windows relative to odor onset. (**D**) Odor concentration classification accuracy of a linear SVM, trained and tested on response vectors from 250 ms sliding windows, using pseudo-populations of 75 cells (100 repeats). Decoding was performed separately for each odor identity, and classification accuracies were then averaged across odors. Dashed line indicates chance level (33.3%). Shading around lines: 95% CI. (**E**) Odor concentration classification accuracy of a linear SVM, trained (x-axis) and tested (y-axis) on response vectors from non-overlapping 500 ms windows, using pseudo-populations of 75 cells, with 100 train/test repeats.

To quantify how identity and concentration contributed to these responses, we performed time-resolved two-way ANOVAs (factors: odor identity and concentration) and measured the fraction of cells significantly modulated by each term. Across time, similar fractions of OB-p and mPFC-p neurons were significantly tuned to odor identity (**Figure 3B**), consistent with both populations encoding which odor was present. By contrast, OB-p neurons were more likely to show significant main effects of concentration, as well as identity × concentration interactions, particularly during and shortly after odor presentation, whereas such effects were rare in mPFC-p neurons (**Figure 3B,C**). Extended metrics of responsiveness and sparseness indicated that, across the three concentrations, overall response sparseness and the fraction of active cells changed only modestly in all three populations (**Figure S5**), suggesting that concentration primarily reshapes response magnitude patterns within a largely stable, sparse ensemble.

We then asked whether these projection-defined ensembles differ in how well they support concentration readout at the population level. We trained linear classifiers on size-matched pseudo-populations to decode odor concentration. OB-p pseudo-populations reached higher concentration classification accuracy and did so earlier in the trial than both mPFC-p and untagged pseudo-populations, whereas mPFC-p ensembles supported only weak concentration decoding that remained close to chance (**Figure 3D,E** and **Figure S5F**). When decoding was restricted to pairwise discriminations between low vs. medium, medium vs. high, and low vs. high concentrations for each odor, concentration decoding was again strongest and most sustained in OB-p ensembles, intermediate in untagged neurons, and weakest in mPFC-p neurons (**Figure S5G**). Together, these analyses indicate a marked specialization of OB-p feedback neurons in encoding odor concentration, whereas mPFC-p projection neurons largely preserve odor identity with comparatively weak concentration dependence.

### Differential encoding of odor valence in OB-p and mPFC-p neurons

Odors can acquire motivational value through learning and experience. Given the role of mPFC in encoding stimulus valence (51–53), we next tested whether odor valence was preferentially encoded in mPFC-p neurons compared to OB-p neurons.

To address this question, we used a Go/NoGo odor discrimination task. We trained mice to lick in response to two odorants previously associated with water reward (CS+1 and CS+2), but to refrain from licking in response to two new, unrewarded odorants (CS-1 and CS-2) (**Figure 4A, B**, **Methods** and **Table 4**). Behavioral performance improved as training progressed, with no significant difference between OB-p and mPFC-p labelled mice (**Figure 4C**; repeated-measures ANOVA within days: F = 4.375, p = 0.0374; a mixed ANOVA within days p = 0.0441, between projections p = 0.767, without a significant interaction p = 0.421), with, on average, 73.7% correct trials (hit and correct rejection) on day 3 and an increase in anticipatory licking in response to the rewarded odorants (**Figure 4C**, and **Figure S6**).

**Figure 4.**
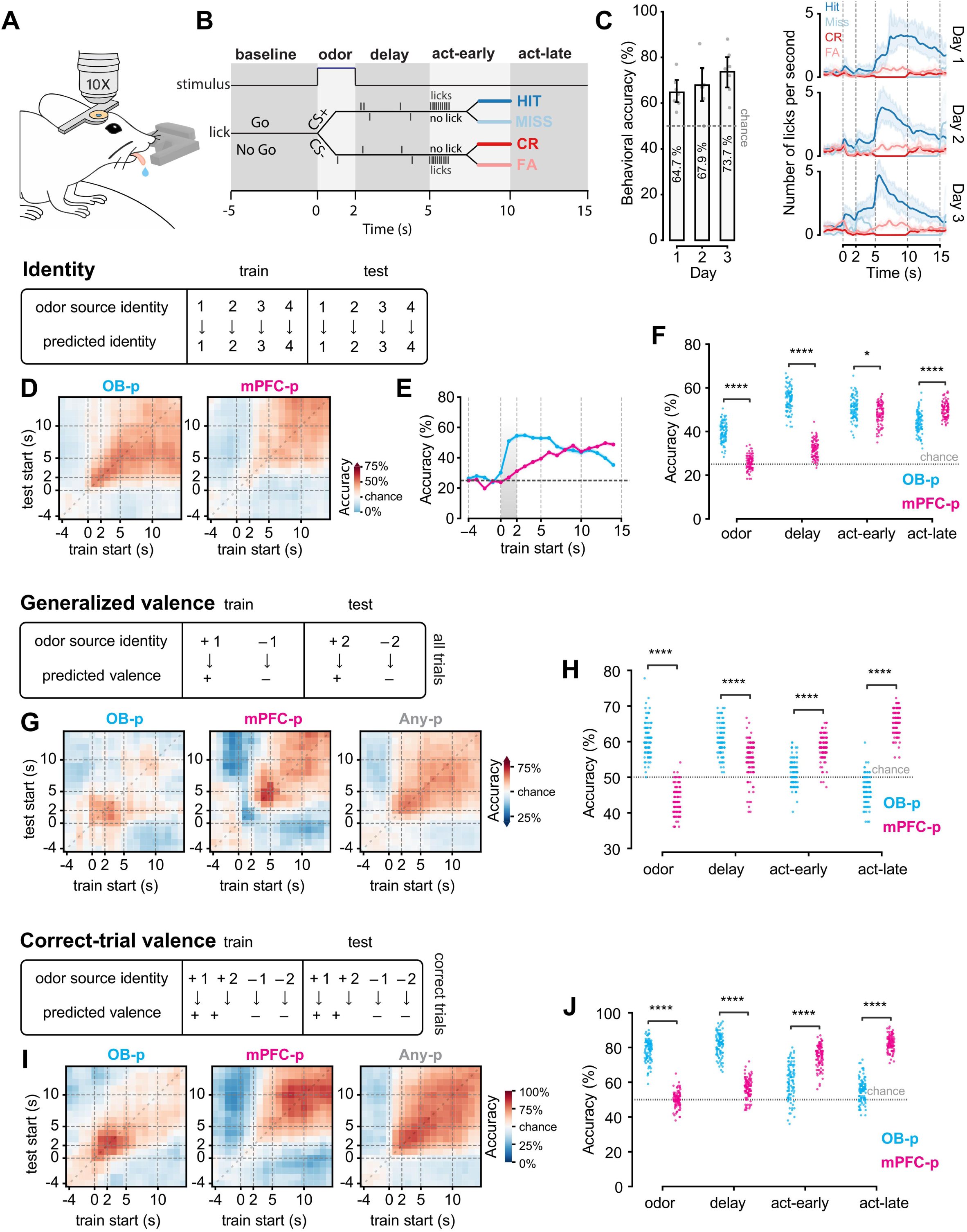
Differential encoding of odor valence in OB-p and mPFC-p neurons. (**A**) Two-photon imaging setup during the odor discrimination task (Go/NoGo). (**B**) Schematic of Go/NoGo task design. Mice learned to lick (Go) for a rewarded odor (CS+1, CS+2) and withhold licking (NoGo) for a non-rewarded odor (CS-1, CS-2). Task phases used for analysis are shown at the top. Trials are divided into four consecutive epochs, relative to odor onset: odor (0-2 s), delay (2-5 s), act-early (5-10 s), act-late (10-15 s). (**C**) Left: Behavioral accuracy across days. Right: Lick rate for each behavioral outcome across days. Shading around lines: 95% CI. Animals performed more accurately and reacted earlier as learning progressed. See also **Figure S6**. (**D**) Dynamics of odor identity decoding using pseudo-populations. Each heatmap shows odor identity classification accuracy of a linear SVM (white: chance level at 25%) using 1-second train (x-axis) and 1-second test (y-axis) non-overlapping windows. Data were averaged over 100 repeats with size-matched pseudo-populations (85 cells). (**E**) Decoding accuracy when pseudo-populations were trained and tested on the same 1-second windows. Dashed line: chance level at 25%. Shaded vertical column: 95% CI. (**F**) Odor identity decoding in expert mice (Day 3) across task phases. Odor identity was classified using a linear SVM with size-matched pseudo-populations (85 cells, sampled 100 times) from projection neurons (colors). Significant differences were found in decoding accuracy between OB-p and mPFC-p pseudo-populations across all task phases. (**G**) Dynamics of generalized valence decoding using pseudo-populations in expert animals (Day 3). Each heatmap shows valence classification accuracy of a linear SVM (white = chance level at 50%) using 1-second train (x-axis) and 1-second test (y-axis) non-overlapping windows. Any-p represents a general PCx population, which is sampled from both projection and untagged neurons. Data were averaged over 100 repeats with size-matched populations (85 sampled cells). (**H**) Generalized valence decoding across task phases using sized-matched (85 cells sampled for 100 repeats) projection pseudo-populations (colors) in expert animals (Day 3). Significant differences were found in decoding accuracy between OB-p and mPFC-p pseudo-populations across all task phases. (**I-J**) Valence decoding restricted to correct trials (hits and correct rejections) in expert animals (Day 3). Days 1-2 were excluded due to imbalanced counts of correct trials across animals to properly construct pseudo-populations. Panels (**I, J**) mirror (**G, H**). See also **Figure S8E** and **G** for generalized valence in correct trials.

Pseudo-population analysis of neural activity from 4 OB-p tagged mice and 3 mPFC-p tagged mice showed that, in expert animals (day 3), odor identity decoding in OB-p pseudo-populations was greater than chance across all 4 task phases (one-sample one-sided t-test with chance = 25%, with Bonferroni corrected p-values across phases: odor = 2.34 × 10⁻^56^, delay = 7.27 × 10⁻^78^, act-early = 1.66 × 10⁻^73^, act-late = 1.2 × 10⁻^63^). Decoding accuracy peaked during early trial phases (delay phase after odor exposure), but declined later in the trial (action) (**Figure 4D, E** and **F**). In contrast, mPFC-p pseudo-populations performed near chance during early trial phases, but decoding accuracy increased later in the trial (one-sample one-sided t-test with chance = 25%, with Bonferroni corrected p-values across phases: odor = 0.0301, delay = 1.42 × 10⁻^32^, act-early = 4.44 × 10⁻^70^, act-late = 4.95 × 10⁻^87^).

To test whether the odor-reward associations were differentially encoded in OB-p versus mPFC-p neurons, we next trained an SVM classifier to predict rewarded (CS+) versus non-rewarded (CS-) odorants based from pseudo-population activity. To exclude confounding information about odor identity from the analysis, we trained the classifier on one CS+/CS-pair, and tested on the other (e.g., train on CS+1 and CS-1 trials, then test on CS+2 and CS-2 trials), with all combinations used (generalized valence, **Figure 4G**, and **Methods**; contrast with **Figure S8A-D** and **I** for unconstrained valence decoding). We found that odor valence could be decoded with high accuracy from a general PCx pseudo-population across all phases (**Figure 4G**, also see **Figure S8H**, one-sample one-sided t-test with chance = 50%, with Bonferroni corrected p-values across phases: odor = 0.00222, delay = 5.31 × 10⁻^31^, act-early = 1.55 × 10⁻^28^, act-late = 1.7 × 10⁻^16^).

Interestingly, we found that OB-p pseudo-populations accurately encoded odor valence during early phases of the trial (odor exposure, delay), but that decoding accuracy decreased to chance level later in the trial (action) (**Figure 4H**; one-sample one-sided t-test with chance = 50%, with Bonferroni corrected p-values across phases: odor = 1.37 × 10⁻^40^, delay = 1.78 × 10⁻^44^, act-early = 0.00141, act-late = 1). In contrast, mPFC-p pseudo-populations encoded odor valence during later but not early phases of the trial (one-sample one-sided t-test with chance = 50%, with Bonferroni corrected p-values across phases: odor = 1, delay = 2.62 × 10⁻^15^, act-early = 1.07 × 10⁻^42^, act-late = 2.42 × 10⁻^68^). We found statistically significant differences in decoding accuracy across all four phases between projection pseudo-populations (two-sided Mann-Whitney U test with Bonferroni corrected p-values across phases: odor = 1.29 × 10⁻^33^, delay = 3.05 × 10⁻^13^, act-early = 1.14 × 10⁻^23^, act-late = 1 × 10⁻^33^).

Similar results were obtained when restricting valence decoding analyses to correct trials, to exclude potential confounds from incorrect behavioral choices (correct trial valence, **Figure 4I** and **J**, see also **Figure S8J**). OB-p pseudo-populations accurately encoded odor valence during early phases of the trial but much less so later (one-sample one-sided t-test with chance = 50%, with Bonferroni corrected p-values across phases: odor = 5.04 × 10⁻^69^, delay = 4.35 × 10⁻^72^, act-early = 4.29 × 10⁻^18^, act-late = 3.04 × 10⁻^12^). In contrast, mPFC-p pseudo-populations encoded odor valence late in correct trials more accurately than early (one-sample one-sided t-test with chance = 50%, with Bonferroni corrected p-values across phases: odor = 1, delay = 1.97 × 10⁻^19^, act-early = 3.89 × 10⁻^63^, act-late = 1.16 × 10⁻^88^).

For all four task phases, these projection pseudo-populations performed differently from one another at decoding valence for correct trials (**Figure 4J**; two-sided Mann-Whitney U test with Bonferroni corrected p-values across phases: odor = 1.09 × 10⁻^33^, delay = 1.46 × 10⁻^33^, act-early = 5.06 × 10⁻^22^, act-late = 1.12 × 10⁻^33^). We also observed significant differences with generalized valence decoding performance in correct trials between projection pseudo-populations during odor and act-early phases (**Figure S8E,G** and **K**; two-sided Mann-Whitney U test with Bonferroni corrected p-values across phases: odor = 4.54 × 10⁻^31^, delay = 0.86, act-early = 2.11 × 10⁻^27^, act-late = 1).

Together, these results reveal a pronounced temporal segregation in the representation of odor identity and valence: OB-p neurons encode odor identity and valence during odor exposure, whereas mPFC-p neurons do so during reward-related behavior.

### Longitudinal tracking reveals greater stability of valence encoding in feedforward projection neurons

Given the prominent role of prefrontal circuits in linking sensory cues to learned outcomes, we next asked whether feedforward mPFC-p neurons maintain more stable valence representations across learning than feedback OB-p neurons. To address this, we longitudinally tracked identified neurons across three Go/NoGo sessions and followed 46 OB-p and 41 mPFC-p neurons in 4 OB-p–tagged and 3 mPFC-p–tagged mice (**Figure 5A, B** and **Figure S10A**). This approach allowed us to quantify learning-dependent changes at the single-cell level and to assess how information is preserved within a fixed ensemble over days.

**Figure 5.**
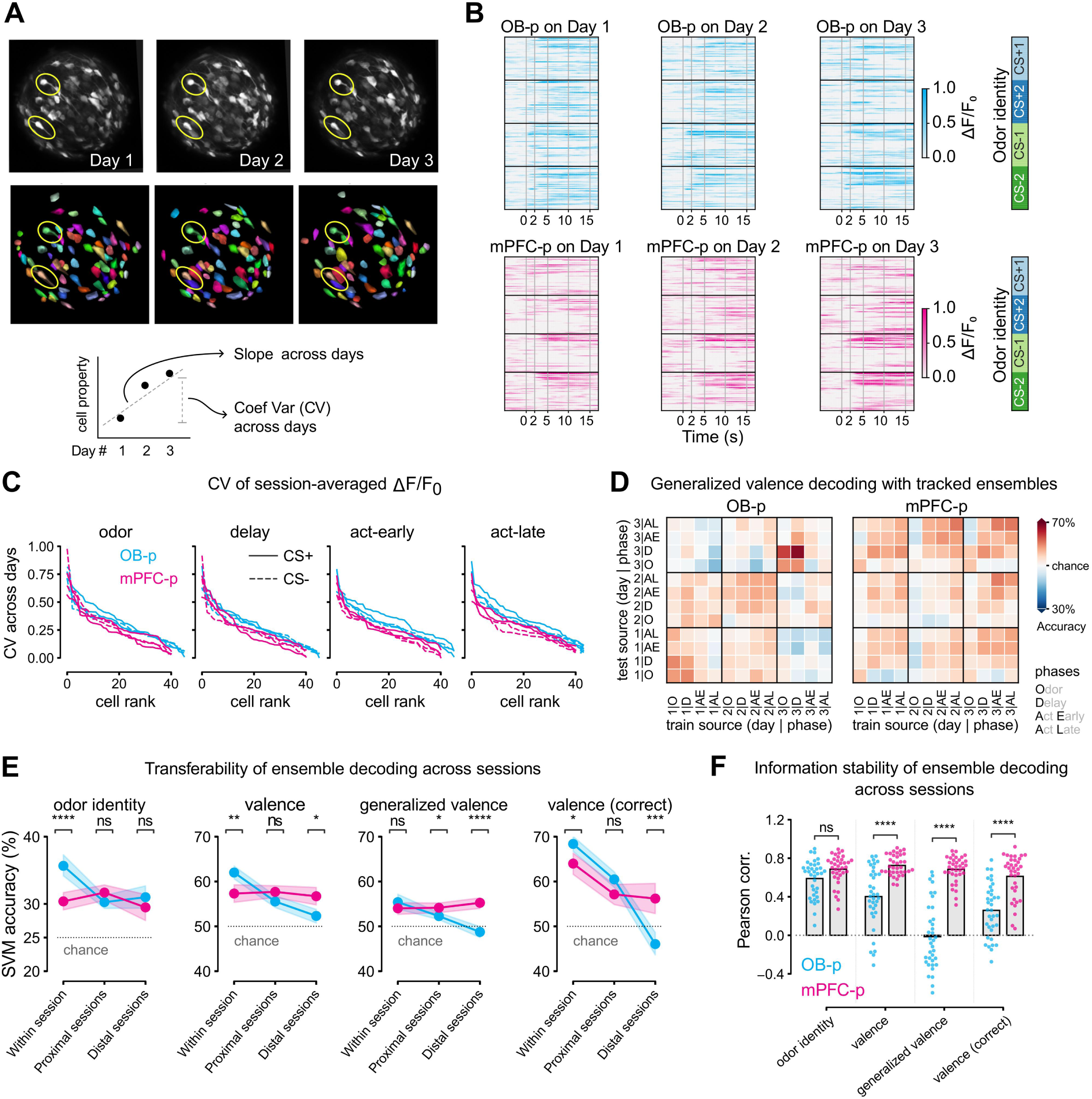
mPFC-projecting ensemble maintain more stable odor valence coding than OB-projecting ensemble. (**A**) Top: Example of tracked cells using ROICaT (top: field-of-view image per session; bottom: segmented cells, same colors indicate same cells; yellow outlines highlight two example tracked cells across days). See **Figure S10** and **Methods** for more details on longitudinal tracking of projection neurons. Bottom: Illustration of session-to-session (restricted to Days 1-3) analyses of single-cell properties for a given tracked neuron. The slope of a linear fit estimates the effects of learning. The coefficient of variation (CV) represents the normalized variability across learning. (**B**) Trial-averaged fluorescence (ΔF/F₀) for all OB-p and mPFC-p neurons tracked across the three Go/NoGo training sessions. Each row represents an individual neuron followed across days, and each column corresponds to a session. The vertical cell ordering is kept the same across odors and days. (**C**) CV across days of averaged fluorescence activity is plotted for OB-p and mPFC-p neurons. Colors: projection; Solid lines: CS+, dashed lines: CS-. Each panel is a different task phase. The CV for each cell is ranked for each panel, stimulus, and projection target separately. See **Results** and **Figure S10F** for statistical comparisons. (**D**) Each heatmap shows the accuracy of generalized valence classification for a linear SVM, (left: OB-p, right: mPFC-p), when trained (x-axis) and tested (y-axis) on different combinations of phases (O: odor, D: delay, AE: act-early, AL: act-late) and days (1, 2, 3). Tracked ensembles for each animal were trained and tested separately, then averaged across animals. Encoding of generalized valence was more transferable across sessions in mPFC-p ensembles than OB-p ensembles. For other decoding schemes, see **Figure S11A**. The data in this panel and the **Figure S11A** were further analyzed in panels (**E, F**) and **Figures S12, S13** for transferability and information stability analyses. (**E**) Accuracy of classification for linear SVM classifiers, trained to decode identity and different types of valences, and tested across sessions. To avoid comparisons between discontinuous task phases (e.g., odor and act-late being grouped together), only adjacent and same-phase groupings were considered (e.g., odor and odor; odor and delay). For train/test on different sessions, “proximal” refers to train/test on consecutive sessions, while “distal” refers to discontinuous sessions (Day 1 and Day 3). mPFC-p ensembles exhibited more transferability in valence decoding than OB-p ensembles. See **Figure S11B** for demonstration of this computation with simulated data, as well as **Figure S12** for additional transferability analyses for different decoding tasks with phase and day difference considerations. (**F**) Following odor identity and valence classification with linear SVM classifiers in **5D**, decoding stability is compared between tracked mPFC-p and OB-p ensembles. Each transfer block consists of how a tracked ensemble SVM trained on one day performs on another day across the different task phases, i.e. each small square 4×4 matrix as illustrated in panel (**D**). Session-to-session pairwise Pearson correlations are shown for each projection ensemble (colors) and selected decoding targets (x-axis). Higher stability for valence information was observed for mPFC-projecting ensembles. See **Figure S11B** for demonstration of this computation with simulated data, and the difference between “transferability” and “stability” used here. See **Figure S13** for analyses of other SVM decoding schemes, as well as other correlation methods in addition to Pearson.

At the single-neuron level, we first examined whether average response magnitudes showed systematic drifts across training. For each tracked cell, we computed the slope of its session-averaged normalized fluorescence across the three Go/NoGo days (**Figure S10A**, bottom; **Figure S10B**). Individual neurons in both populations increased or decreased their response magnitudes over sessions, but there was no significant difference in the average slope between OB-p and mPFC-p neurons (**Figure S10D**; mixed ANOVA on slopes within combinations of odor valence and task phase: main effect of combination p = 0.110; main effect of projection target p = 0.360; interaction p = 0.00310). We similarly quantified changes in lifetime sparseness across sessions and did not observe a clear effect of projection target (**Figure S10E**; mixed ANOVA on sparseness slopes within task phases: main effect of task phase p = 0.0491; main effect of projection target p = 0.364; interaction p = 0.765). Thus, on average, response amplitudes and selectivity did not exhibit strong monotonic drifts that differed between OB-p and mPFC-p neurons.

Although monotonic changes were similar across projections, we did detect differences in response variability. We quantified, for each tracked neuron, the coefficient of variation (CV) of its session-averaged normalized fluorescence (**Figure 5A**, bottom) and compared matched-ranked OB-p and mPFC-p neurons. OB-p neurons typically showed higher fluorescence CV across days than mPFC-p neurons (**Figure 5C, D**). A mixed ANOVA on CV within combinations of odors and task phases revealed a significant effect of projection target (p = 0.0127) and of odor–phase combination (p = 0.0372), without a significant interaction (p = 0.43). More detailed post-hoc comparisons across specific conditions did not reach significance after correcting for multiple Mann–Whitney U tests (**Figure S10F**), but overall suggest greater trial-to-trial and session-to-session variability in OB-p neurons.

We next asked how these differences manifest at the population level by using classifier-based analyses on the tracked ensembles. Longitudinal registration enabled us to evaluate information transfer within the same encoding space, by training a classifier on one day and testing it on another. When decoding odor identity, transferability across dissimilar sessions was comparable between OB-p and mPFC-p ensembles (**Figure 5E**, first panel; two-sided Mann–Whitney U tests with Bonferroni correction across session-pair transfers: within = 4.59 × 10^-5^, proximal = 0.194, distal = 0.686). In contrast, across multiple valence decoding schemes, OB-p ensembles showed a more pronounced decline in performance when classifiers were transferred across days, whereas mPFC-p ensembles maintained higher valence decoding accuracy, particularly for generalized valence across distal sessions (**Figure 5E** and **Figure S11**; two-sided Mann–Whitney U tests with Bonferroni correction for generalized valence: within = 0.774, proximal = 0.0356, distal = 5.41 × 10^⁻6^).

To further probe the stability of temporal coding, we correlated the session-transfer decoding time courses across different days. These analyses revealed that the temporal structure of valence coding was consistently more stable in mPFC-p than in OB-p ensembles (**Figure 5F** and **Figure S12B**; two-sided Mann–Whitney U tests with Bonferroni correction across four decoding schemes: odor identity p = 0.755, valence p = 6.24 × 10^-6^, generalized valence p = 2.89 × 10^-11^, valence for correct trials p = 2.50 × 10^-6^; see also **Figure S13** for similar results using alternative correlation methods).

Together, longitudinal tracking of identified neurons indicates that, although single-cell response magnitudes and sparseness change similarly across learning in OB-p and mPFC-p populations, valence representations are more stable and more transferable over days in feedforward mPFC-p ensembles than in feedback OB-p neurons.

## Discussion

In this study, we employed two-photon calcium imaging and viral labeling to investigate differences in odor coding between OB-projecting (OB-p) and mPFC-projecting (mPFC-p) neurons in the piriform cortex (PCx). By analyzing single-neuron and population-level properties, we identified differences in information capacity, temporal dynamics, and plasticity. These findings reveal differences in sensory representation between feedforward and feedback projection neurons in the olfactory cortex, with implications for understanding the transformation of sensory representations across hierarchical processing pathways.

### Temporal dynamics of projection neurons align with the hierarchical organization of olfactory processing

We found that OB-p and mPFC-p ensembles exhibited distinct temporal dynamics in response to passive odor presentation and during an associative olfactory learning task. Using a transformer model, we were able to classify projection targets based on neural activity. Analysis of the model’s attention weights suggested differences in the temporal dynamics of these projection neurons, prompting us to perform detailed analyses of their functional properties and information capacity. Specifically, we found that OB-p neurons responded to odorants with a shorter latency. Using pseudo-population analyses with linear SVM classifiers, we also found that OB-p ensembles encoded odor identity and concentration earlier in the trial than mPFC-p ensembles. Finally, in expert animals trained on the Go/NoGo odor discrimination task, OB-p neurons encoded identity and reward associations earlier in the trial than mPFC-p neurons. These observations suggest that OB-p neurons play an important role in sensory processing within the first few seconds of odor exposure, consistent with a role in the real-time updating of olfactory bulb mitral and tufted cell outputs (30,32,50,54–56). Differences in the dynamics of odor information encoding may reflect differential connectivity with olfactory bulb mitral and tufted cells as well as within PCx, such as differences in the dependence on recurrent circuitry to drive excitation (31,57–62). In addition, differential dynamics could result from the activity of feedback loops between PCx and OB and mPFC, respectively (63–66). Differential connectivity may thus explain why OB-p and mPFC-p neurons preferentially represent odor information across different task phases, including odor ON and OFF responses (**Figure 6**; Wilson 1998).

**Figure 6.**
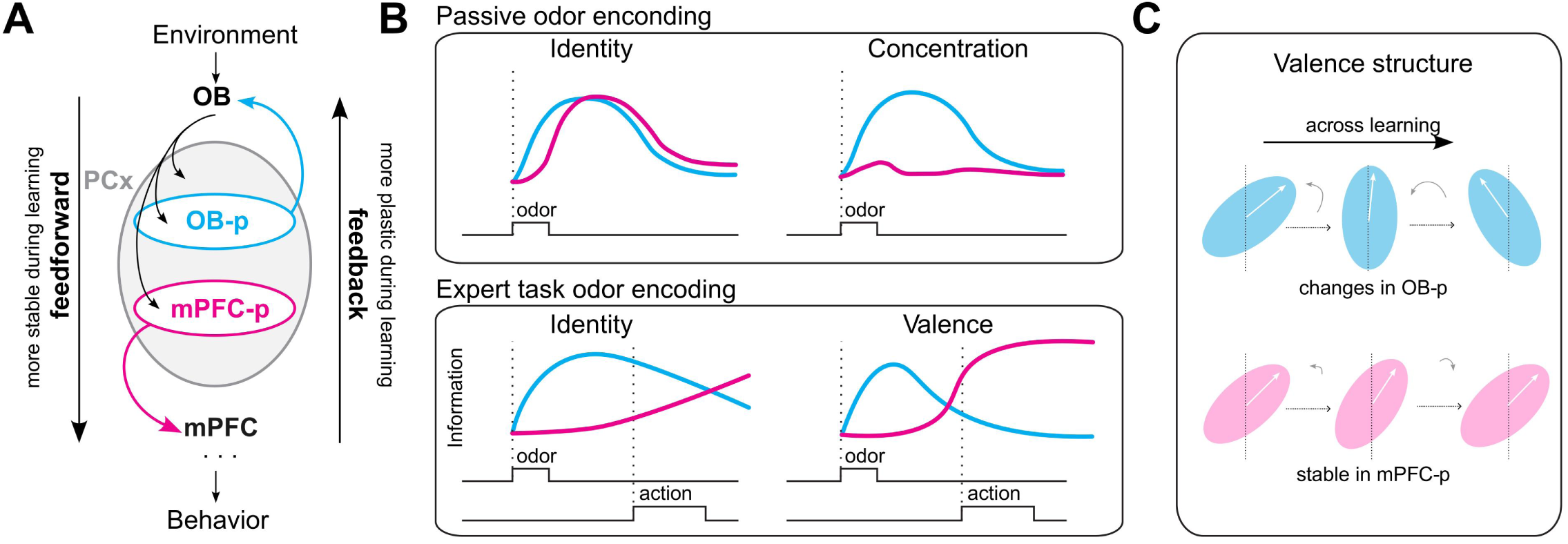
Olfactory information flows through temporally distinct circuits: fast, plastic feedback, and slow, stable feedforward pathways. **(A)** Schematic illustrating the two projection-defined pathways studied here. OB-p neurons send feedback projections from PCx to the olfactory bulb whereas mPFC-p neurons send feedforward projections to medial prefrontal cortex. **(B)** During odor discrimination in expert animals, OB-p ensembles encode odor identity earlier than mPFC-p ensembles, while OB-p ensembles more robustly encode odor concentration. In contrast, mPFC-p ensembles encode generalized and behavior-relevant valence at later phases in the task, which are typically associated with animal behaviors or cognitive processes. **(C)** Summary of longitudinal analyses. Across learning, the structure of valence information is more stable in mPFC-p ensembles than in OB-p ensembles, reflecting a division of labor in which feedback pathways provide rapidly updated sensory information and feedforward pathways maintain stable higher-order representations.

### Functional specializations of projection neurons align with those of their target areas

During an associative olfactory learning task, OB-p and mPFC-p ensembles exhibited distinct temporal dynamics that persisted beyond odor detection and continued throughout the ensuing behavioral response. OB-p neurons encoded odor identity early during odor exposure, while mPFC-p neurons encoded odor identity later in the trial, when reward-related behaviors took place. Similarly, both subpopulations encoded odor-reward associations, but OB-p ensembles did so earlier than mPFC-p ensembles. Notably, these differences increased over the course of learning. The encoding of odor-reward associations in OB-p neurons suggests that OB circuits not only perform basic signal processing functions such as signal normalization but also have access to higher-order olfactory information. This finding extends prior observations obtained from recording axonal and somatic neural activity in the OB (68–71). In contrast, feedforward mPFC-p ensembles are preferentially engaged during the animal’s behavioral response to the odor stimulus. The later engagement of mPFC-p neurons aligns with the well-established role of mPFC in higher-order cognitive processes (53,72–74). The finding that feedback projections neurons in PCx more rapidly encode odor information than feedforward neurons may reflect a general principle of the hierarchical processing of sensory signals. For example, in the barrel cortex, cortico-thalamic (CT) neurons exhibit shorter spiking latencies than cortico-cortical (CC) neurons, which include neurons projecting to higher-order cortical areas (1–6,75). Taken together, our findings support a model where the short-term temporal dynamics of sensory cortex neurons align with the functional demands of their projection targets.

### Feedback pathways exhibit greater modulation by learning than feedforward pathways

Tracking individual neurons across behavioral sessions allowed us to investigate the effect of learning on the functional properties of OB-p and mPFC-p ensembles. At the single-neuron level, average changes in response magnitude and lifetime sparseness across days were similar between the two projection types, but OB-p neurons exhibited higher variability in their fluorescence across sessions. These observations suggest that, during learning, individual OB-p neurons show more labile response patterns, whereas mPFC-p neurons display more stable single-cell response profiles over time.

Beyond single-cell changes, our longitudinal decoding analyses further highlighted a dissociation between identity and valence stability across projection-defined ensembles. When classifiers were trained on one session and tested on another, odor identity decoding transferred similarly across days in OB-p and mPFC-p populations, indicating that basic identity representations remained relatively stable in both pathways. By contrast, multiple valence-related decoding schemes, including generalized valence across odors and valence restricted to correct trials, showed a more pronounced decline in transferability for OB-p ensembles than for mPFC-p ensembles. Together, these results suggest that feedforward pathways carrying higher-order olfactory information such as odor valence exhibit greater stability, whereas feedback pathways exhibit greater plasticity. This selectivity in the stability of information routing may more effectively facilitate the updating of network properties with context-relevant information during learning (12,76,77).

### Projection-specific division of labor in piriform cortex

Taken together, our findings support a model in which projection target defines a major organizing axis of odor coding in piriform cortex. OB-p feedback neurons respond with short latency, exhibit strong concentration-dependent modulation, and show substantial learning-related changes in valence-related population structure. In contrast, mPFC-p feedforward neurons engage later in the trial, display weaker concentration dependence, and maintain more stable valence representations across extended training. The untagged population often showed intermediate properties, suggesting that projection-specific biases are superimposed onto a broader piriform representation, rather than forming completely segregated, projection-specific codes. These complementary specializations (summarized in **Figure 6**) are consistent with a division of labor in which OB-p ensembles provide rapidly updated, intensity-rich sensory signals to modulate early olfactory processing in the bulb, whereas mPFC-p ensembles convey more stable, concentration-tolerant representations of odor identity and valence to higher-order cortical circuits involved in decision making and outcome evaluation.

The present work focuses on two projection-defined populations, yet piriform cortex sends outputs to multiple downstream targets, including orbitofrontal cortex, amygdala, entorhinal cortex, and hypothalamus (28,31). An important next step will be to determine whether these additional projection pathways form a broader family of parallel channels with distinct temporal dynamics, concentration sensitivity, and learning rules, or whether OB-p and mPFC-p neurons represent two extremes along a continuum of projection-specific coding strategies. Combining the type of projection-specific imaging used here with cell-type–specific perturbations and chronic recordings over longer time scales will be essential to test how these pathways causally contribute to odor-guided decisions and the updating of odor value.

More generally, our results raise the possibility that a similar division of labor between fast, plastic feedback pathways and slower, more stable feedforward pathways may be a common organizing principle across sensory cortices. In such a framework, feedback projections would continuously reshape early sensory representations based on current context, internal state, and recent experience, while feedforward projections maintain stable, behaviorally relevant readouts for downstream decision circuits. Future work examining projection-specific coding across different behavioral states and task conditions will be important to determine how flexible this division of labor is, and whether it can be dynamically reconfigured to match the demands of the environment.

## Data and code availability

Datasets are converted to the NWB format (NeurodataWithoutBorders) and stored in the Dandi Archive: https://dandiarchive.org/dandiset/000785/. All data will be made publicly available upon publication.

A repository to explore the data is currently available at: https://gitlab.com/fleischmann-lab/calcium-imaging/projection-difference/

The code repository that contains full analysis scripts and notebooks will be made available on the lab GitLab repository upon publication: https://gitlab.com/fleischmann-lab/

Parts of this research were conducted using computational resources and services at the Center for Computation and Visualization, Brown University.

## Acknowledgements

We thank D. Sheinberg, A. Schaefer, Y. Ma, T. Serre, J. Ritt, O. McKissick, the U19 Osmonaut consortium, and members of the Fleischmann Lab for valuable feedback, discussions, and technical assistance. We are grateful to J. Zapata for initial training on GRIN lens implant procedures and to J. Murphy for his help in designing behavioral rigs. We thank A. Pierré for software support. Research in the Fleischmann Lab was supported by grants from the NIH (NIDCD R01DC017437 and 1U19NS112953-01) and by the Robert J. and Nancy D. Carney Institute for Brain Science. Research in the Franks Lab was supported by grants from the NIH (DC015525) and research in both the Fleischmann and Franks Labs was supported by a grant from the NSF (DC016782). Research in the Dyer Lab was supported by grants from the NIH (1R01EB029852) and the NSF (CAREER IIS-2146072), as well as generous gifts from the CIFAR Learning in Machines & Brains Program and The Hypothesis Fund. Carney Institute computational resources used in this work were supported by the NIH Office of the Director (S10OD025181).

## Author contributions

*SD*, *AF*, and *KMF* conceptualized the study with input from all authors. *SD* conducted all experiments and, together with *TP*, curated the data. Formal analyses were performed by *TP* and *SD*. *AA* performed the transformer analysis with help from *JX*. *SS* performed the reliability analysis. *TP* and *SD* developed the software and generated the visualizations. *AS*, *CN*, and *TF* assisted with GRIN lenses. Project administration was overseen by *AF*, and funding was acquired by *AF*, *ED*, and *KMF*. Supervision and scientific guidance were provided by *AF*, *ED*, and *KMF*. The manuscript was written by *SD*, *TP*, *MS*, *KMF*, and *AF* with input from all authors.

## Competing interests

The authors declare no competing interests.

## Declaration of generative AI and AI-assisted technologies

During the preparation of this work, the authors used ChatGPT (OpenAI) to assist with text formatting and reviewing code. After using this tool, the authors reviewed and edited the content as needed and took full responsibility for the content of the publication.

## Methods

**Table 1:** Key resources.

| Reagent type or resource | Designation | Source of reference | Identifier | Additional information |
| --- | --- | --- | --- | --- |
| Strain (M. musculus) | C57BL/6J | Jackson Laboratory | Stock #: 000664<br>RRID:IMSR_JAX:000664 |  |
| Strain (M. musculus) | Ai14/ B6.Cg-Gt(ROSA)26Sortm14(CAG-tdTomato) Hze/J | Jackson Laboratory | Stock #: 007914<br>RRID:IMSR_JAX:007914 |  |
| Genetic reagent (Adeno-Associated Virus) | pGP-AAV-syn-jGCaMP7f-WPRE | Addgene | 104488-AAV1 | AAV1 serotype, diluted 1:3 in sterile 1× PBS |
| Genetic reagent (Adeno-Associated Virus) | pENN.AAV.hSyn.Cre.WPRE.hGH | Addgene | 105553-AAVrg | AAV-retro, used at stock concentration (no further dilution) |
| Software, algorithm | Suite2p | https://github.com/MouseLand/suite2p |  |  |
| Software, algorithm | ROICAT | https://github.com/RichieHakim/ROICaT |  |  |
| Software, algorithm | Manual refinement GUI for tracked neurons | TBD |  |  |
| Software, algorithm | Python, scikit-learn |  |  |  |
| Software, algorithm | Fiji/ImageJ |  |  |  |
|  | DANDI Archive (embargoed) | https://dandiarchive.org/dandiset/000785 |  |  |
|  | Demonstrative code repository with DANDI | https://gitlab.com/fleischmann-lab/calcium-imaging/projection-difference |  |  |
|  | Main analysis codes [TBD] |  |  |  |

### Experimental design and subjects

All experiments and surgical procedures were conducted in accordance with the Guide for the Care and Use of Laboratory Animals (National Institutes of Health) and were approved by the Institutional Animal Care and Use Committee (IACUC) at Brown University (protocol numbers 21-03-0004 and 24-03-0004). Mice were housed in a temperature- and humidity-controlled facility under a 12-h light/dark cycle with food and water available ad libitum (unless otherwise noted). A total of 18 Ai14 (B6.Cg-Gt(ROSA)26Sor<tm14(CAG-tdTomato)>Hze/J × C57BL/6J) adult mice (both males and females, 8-12 weeks old) were used for this study. Of these, 9 mice received retrograde AAV injections into the olfactory bulb (OB) and are referred to as “OB-p” mice; 8 received injections into the infralimbic/prelimbic cortex (mPFC) and are referred to as “mPFC-p” mice. For imaging experiments, mice were selected based on the robustness of viral expression and optical clarity post-surgery. Mice were housed singly after surgery to protect the head-mounted gradient index (GRIN) lens.

#### Group assignments and exclusion criteria

Passive Odor Sessions: 16 mice underwent passive odor exposure sessions to characterize odor-evoked responses (identity and/or concentration dataset). Go/NoGo Training: A subset of these mice (4 OB-p and 3 mPFC-p) proceeded to the head-fixed Go/NoGo paradigm. Exclusions: Mice were excluded from final analysis if (1) the GRIN lens placement was inaccurate (verified post hoc by histology), (2) there was negligible expression of jGCaMP7f or tdTomato in the anterior piriform cortex (aPCx), or (3) the animal failed to learn basic licking behavior during the habituation phase.

### Stereotaxic surgeries

Mice were anesthetized with isoflurane (induction at 3%, maintenance at 1-2% in 1 L/min O2) and placed in a stereotaxic frame (David Kopf Instruments). Ophthalmic ointment (Puralube) was applied to prevent corneal desiccation, and body temperature was maintained at 37 °C using a feedback-controlled heating pad (Harvard Apparatus). Extended-release buprenorphine (0.05-0.1 mg/kg, subcutaneous) was administered at least 15 min before incision for analgesia. The scalp was shaved, and the incision area was disinfected with alternating wipes of 70% ethanol and povidone-iodine.

A small craniotomy (∼1 mm diameter) was drilled above the right aPCx at the following coordinates (mm) relative to bregma: ML = 3.7, AP = 0.3. We used a glass micropipette (pulled with a Sutter Micropipette Puller, tip diameter ∼20-30 µm) to deliver pGP-AAV-syn-jGCaMP7f-WPRE at a rate of 100 nL/min. The pipette was slowly advanced to three depths (DV ≈ −3.6, −3.7, and −3.9 mm) to deliver a total volume of ∼950-1100 nL across the three sites. After each injection, the pipette was left in place for 5 min to allow viral diffusion, then retracted at 200 µm/min. A second craniotomy was performed either above the olfactory bulb (OB) or the infralimbic/prelimbic regions of medial prefrontal cortex (mPFC); OB target: ML = 1.0, AP = 4.5, at depths DV = −0.8 and −0.5 mm; mPFC target: ML = 0.4, AP = 1.65, at depths DV = −2.05 and −1.80 mm. At each site, pENN.AAV.hSyn.Cre.WPRE.hGH (AAV-retro) was injected at 100 nL/min for a total of 200-300 nL. After the injection, the pipette was again left in place for 5 min and then retracted at 200 µm/min.

Following the final viral injection, we waited ∼30 min to facilitate initial viral absorption. Next, an aberration corrected 0.5mm diameter gradient index (GRIN, (41), based on a NEM-050-25-10-860-S-2.0p, GRINTech) lens or 0.6mm GRIN lens (NEM-060-25-10-920-S-1.5p, GRINTech) was slowly lowered over the aPCx craniotomy at a rate of 100 µm/min until reaching a depth of ∼−3.5 mm DV (measured from the cortical surface). Excess cerebrospinal fluid or blood was gently wicked away with sterile cotton swabs during the descent. The lens was fixed to the skull with Metabond adhesive cement (Parkell).

A custom aluminum head bar was then placed posterior to the lens and cemented in place using dental acrylic (Pi-ku-plast HP 36, Bredent). Finally, the lens was protected with a silicone elastomer plug (Kwik-Sil, World Precision Instruments). Mice recovered on a warming pad until fully awake and were then returned to their home cage.

Mice were monitored daily for 7 days post-surgery to ensure normal recovery and weight maintenance. For analgesia, a second dose of extended-release buprenorphine (0.05-0.1 mg/kg, subcutaneous) was administered on postoperative Day 3 if needed. Mice were single-housed to protect the head-mounted implant. We allowed 4-6 weeks for viral expression before any *in vivo* imaging experiments. By this time, robust jGCaMP7f fluorescence was typically visible through the GRIN lens.

### Histology and imaging site confirmation

At the conclusion of imaging experiments, mice were deeply anesthetized with an intraperitoneal injection of 2.5% tribromoethanol (Avertin) and perfused transcardially with cold 1x phosphate-buffered saline (PBS), followed by 4% paraformaldehyde (PFA) in PBS. The skull was post-fixed overnight at 4 °C in 4% PFA, after which the brain was dissected out and placed in 4% PFA for an additional 12-24 h.

Fixed brains were rinsed in PBS and embedded in 4% agarose for vibratome sectioning (Leica VT1000 S). Coronal sections of 100 µm thickness were collected across the anterior piriform cortex (aPCx), including the area of the GRIN lens tract and the retrograde injection sites in either the olfactory bulb (OB) or the medial prefrontal cortex (mPFC).

To visualize overall cytoarchitecture, free-floating sections were incubated overnight at 4 °C in NeuroTrace 640/660(1:1000 in 0.1% Triton X-100/PBS). Sections were then washed three times in PBS (15 min each) and mounted onto glass slides using Vectashield Plus mounting medium (Vector Laboratories).

Mounted sections were imaged using a Nikon A1R laser-scanning confocal microscope (Nikon Instruments) equipped with a 10X or 20X objective. Z-stacks were typically collected from 50-130µm thickness with a 5 µm step size to capture the full extent of the injection site and lens track. All confocal settings (laser power, PMT gain) were kept consistent within an experiment to allow for reliable comparison of fluorescence signals.

All Z-stack confocal images were processed in Fiji (ImageJ). Briefly, we used the maximum-intensity projection for initial visualization, followed by background subtraction and brightness/contrast adjustment if needed. For any quantitative ROI-based analyses, we maintained identical settings across all sections from a given experiment.

#### Viral expression

Confocal images were examined to confirm robust tdTomato expression in layer II-III neurons of the aPCx, indicative of projection-specific labeling (OB-p or mPFC-p). OB and mPFC injection sites were similarly inspected for retrograde labeling and to check for any off-target viral spread. (**Supplement Figure S1**)

#### GRIN lens track

We identified the lens track by the absence of tissue and possible gliosis along the dorsoventral axis. Sections containing the lens tract were aligned with the Paxinos and Franklin mouse brain atlas to confirm that the lens tip was positioned just above (∼50-100µm) the targeted aPCx. (**Supplement Figure S1**)

### Animal behavior

#### Water restriction

Mice were water restricted prior to the start of behavioral experiments to motivate licking for water rewards. They received 1 h of controlled water access per day in their home cage. Water restriction began at least 4 days before experiments. Mice were weighed daily throughout the restriction period to ensure they maintained at least 85% of their baseline body weight. If a mouse fell below this threshold, water was provided (up to 2 mL as needed) until its weight recovered.

#### Passive odor exposure

Mice were gradually habituated to being head-fixed in the imaging rig atop a custom 3D-printed wheel for up to 30 min/day for at least 3 days prior to experiment. Odors were delivered through a 16-channel olfactometer (Automate Scientific) at a 1 L/min total flow rate, controlled by a Teensy 3.6 microcontroller. Each odorant or blank (mineral oil, air) was presented in a 22 s trial: 5 s baseline, 2 s odor pulse, and 15 s post-odor interval. Trials were separated by a 10 s inter-trial interval to allow adequate clearing of residual odors (a vacuum port in the imaging box helped evacuate odors).

During passive exposure sessions, mice were head-fixed but did not receive any water rewards. Each odor stimulus was presented 8-10 times in a pseudo-randomized sequence that prevented consecutive repeats of the same odor. Mice typically underwent ∼1 h of passive odor exposure, during which two-photon imaging was performed.

We used 8 monomolecular odorants at a single concentration (0.001-0.1% v/v) for our “identity” panel, and 3 odorants at 3 concentrations each (0.001, 0.01, 0.1% v/v) for our “concentration” panel. Exact odorants, suppliers, and concentrations are listed in **Table 2, 3 and 4**. All odorants were diluted in mineral oil.

**Table 2:** Identity and 30Hz fast imaging dataset.

| Stim name | Pubchem id | Stim types | Stim description | Chemical dilution type | Chemical concentration | Chemical concentration unit | Chemical solvent | Provider |
| --- | --- | --- | --- | --- | --- | --- | --- | --- |
| Mineral Oil | NA | control | Control odor stimulus | NA | NA | NA | NA | Sigma |
| Acetophenone | 7410 | odor | Odor stimulus #1 | vol/vol | 0.01 | % | Mineral Oil | Sigma |
| Amylamine | 8060 | odor | Odor stimulus #2 | vol/vol | 0.01 | % | Mineral Oil | Sigma |
| Butyl Acetate | 31272 | odor | Odor stimulus #3 | vol/vol | 0.01 | % | Mineral Oil | Sigma |
| Ethyl Hexanoate | 31265 | odor | Odor stimulus #4 | vol/vol | 0.01 | % | Mineral Oil | Sigma |
| 2-isobutyl-3-methoxypyrazine | 32594 | odor | Odor stimulus #5 | vol/vol | 0.01 | % | Mineral Oil | Sigma |
| β-Ionone | 638014 | odor | Odor stimulus #6 | vol/vol | 0.1 | % | Mineral Oil | Sigma |
| 2,3-Pentanedione | 11747 | odor | Odor stimulus #7 | vol/vol | 0.001 | % | Mineral Oil | Sigma |
| Valeric Acid | 7991 | odor | Odor stimulus #8 | vol/vol | 0.001 | % | Mineral Oil | Sigma |
| Air | NA | control | Control pure air stimulus | NA | NA | NA | NA | NA |

**Table 3:** Concentration dataset.

| Stim name | Pubchem id | Stim types | Chemical dilution type | Chemical concentration | Chemical concentration unit | Chemical solvent | Provider |
| --- | --- | --- | --- | --- | --- | --- | --- |
| Mineral Oil | NA | control | NA | NA | NA | NA | Sigma |
| Acetophenone | 7410 | odor, A, low | vol/vol | 0.001 | % | Mineral Oil | Sigma |
| Acetophenone | 7410 | odor, A, medium | vol/vol | 0.01 | % | Mineral Oil | Sigma |
| Acetophenone | 7410 | odor, A, high | vol/vol | 0.1 | % | Mineral Oil | Sigma |
| Butyl Acetate | 31272 | odor, B, low | vol/vol | 0.001 | % | Mineral Oil | Sigma |
| Butyl Acetate | 31272 | odor, B, medium | vol/vol | 0.01 | % | Mineral Oil | Sigma |
| Butyl Acetate | 31272 | odor, B, high | vol/vol | 0.1 | % | Mineral Oil | Sigma |
| Ethyl Hexanoate | 31265 | odor, C, low | vol/vol | 0.001 | % | Mineral Oil | Sigma |
| Ethyl Hexanoate | 31265 | odor, C, medium | vol/vol | 0.01 | % | Mineral Oil | Sigma |
| Ethyl Hexanoate | 31265 | odor, C, high | vol/vol | 0.1 | % | Mineral Oil | Sigma |

**Table 4:** Go/NoGo (GNG) dataset.

| Stim name | Pubchem id | Stim types | Stim description | Chemical dilution type | Chemical concentration | Chemical concentration unit | Chemical solvent | Provider |
| --- | --- | --- | --- | --- | --- | --- | --- | --- |
| Ethyl Hexanoate | 31265 | odor, CS+1 | odor stimulus, CS associated with reward, during task | vol/vol | 0.01 | % | Mineral Oil | Sigma |
| β-Ionone | 638014 | odor, CS+2 | odor stimulus, CS associated with reward, during task | vol/vol | 0.1 | % | Mineral Oil | Sigma |
| Amylamine | 8060 | odor, CS-1 | odor stimulus, CS NOT associated with reward, during task | vol/vol | 0.01 | % | Mineral Oil | Sigma |
| Butyl Acetate | 31272 | odor, CS-2 | odor stimulus, CS NOT associated with reward, during task | vol/vol | 0.01 | % | Mineral Oil | Sigma |

#### Go/NoGo odor discrimination task

For Go/NoGo training, a custom 3D-printed mouthpiece (Sandworks, LLC) was positioned in front of the mouse’s snout, equipped with an optical lickometer. A photogate sensor detected each lick, and a stainless-steel spout delivered water rewards via a solenoid valve (controlled by an Arduino Mega2560). An LED mounted in the mouse’s visual field served as a trial-start cue.

Mice underwent pre-training where only one CS+ odor was presented (∼80-100 trials). During this phase, a single 2 s odor stimulus (CS+) was always followed by a water reward (2 µL) after a 3 s delay. Mice typically reached ≥85% hit rate (i.e., licking within 2 s of reward onset) within 1-2 sessions, at which point they progressed to the full Go/NoGo task. We refer to this as “passive day” (Day 0). The days where the animals performed the Go/NoGo task are referred to as “task days/sessions” (Day 1, 2, …).

In the full Go/NoGo task, mice were presented with 4 odors: 2 CS+ (rewarded) and 2 CS− (unrewarded). Each trial consisted of: Cue Onset: A 100 ms LED blink signaled the start of the trial. Baseline (5 s): No odor, no reward possible. Odor (2 s): One of the 4 odors was delivered. Delay (3 s): Mice could not receive reward yet, but could lick freely, and such licks were not considered a response. Outcome/Action Window: Water reward (2 μL) was delivered only on CS+ trials if the mouse licked. Licking on CS− trials was recorded as a false alarm. Inter-Trial Interval (10 s): The olfactometer lines were flushed with clean air to reduce odor carryover.

Approximately 20-30 trials per odor (total ∼80-120 trials/day) were completed for up to 3 consecutive days of training.

We scored each trial as one of the following:

- Hit: Licking within the reward window for a CS+ odor.
- Miss: Failure to lick during the reward window for a CS+ odor.
- False Alarm (FA): Licking during the first 5 seconds of the reward window on a CS− trial.
- Correct Rejection (CR): No licks during the first 5 seconds of the reward window on a CS− trial. Trials where excessive movement or imaging artifacts occurred were excluded from analysis (< 5% of trials). Mice that failed to reach ≥60% correct after 3 days were considered poor learners and excluded from final analyses.

To facilitate neural analyses for action phases (act-early: 0-5 seconds after delay; act-late: 5-10 seconds after delay) and because we did not analyze more than 10 seconds after the delay period, we also relabeled Go trials as hit trials only if the animals hit and received rewards within the first 10 seconds after the delay period.

### Head-fixed two-photon imaging

Two-photon imaging was performed using an Ultima Investigator DL laser scanning microscope (Bruker Nano, Middleton, WI, USA) equipped with an 8 kHz resonant galvanometer and high-speed optics set. Laser excitation was provided by a Chameleon Discovery NX Ti:Sapphire laser (Coherent, Santa Clara, CA, USA) tuned to 960 nm for excitation of both jGCaMP7f and tdTomato. Emitted fluorescence was split by a primary dichroic mirror and detected by dual GaAsP PMTs (Hamamatsu H10770). A secondary beam splitter and emission filters (ET525/70m-2p for GCaMP; ET595/50m-2p for tdTomato) isolated the respective channels.

We used a Nikon 10× Plan Apochromat Lambda objective (0.45 NA, 4.0 mm WD) for two-photon imaging through the GRIN lens. Typical laser power at the sample ranged from 90-150 mW (GRIN lens focal plane), adjusted per mouse to maximize signal-to-noise while minimizing photobleaching and phototoxicity. In some experiments (the “fast_Hz” dataset), we imaged a single optical plane at 512×512 pixels with a frame rate of 30 Hz. This allowed us to capture high temporal resolution activity. In all other experiments, we acquired 3 planes simultaneously using a piezo motor to step the objective ∼80 µm between planes. This yielded an effective frame rate of 4.53 Hz per plane (512×512 pixels). The planes were chosen to span layers II-III of the aPCx, typically 100-500 μm below the lens surface.

After positioning the mouse under the objective, the focal plane was slowly advanced until we visualized the top of the GRIN lens. We then moved 100-500 μm dorsally from the lens surface in 10-20 μm increments to find regions with robust jGCaMP7f expression. Once a suitable field of view (FOV) was located—typically ∼400×400 µm in the brain tissue (depending on the effective magnification)—we fine-tuned laser power to achieve optimal imaging contrast for each plane. A typical imaging session lasted 1.0-1.5 h, during which mice performed either passive odor exposure or a Go/NoGo discrimination task (see §5). Short breaks (1-2 min) were interspersed every ∼15-20 min to relieve potential stress in the animal. After each session, the mouse was returned to its home cage.

For each experiment, raw image frames for both GCaMP7f and tdTomato channels were recorded onto a local solid-state drive via PrairieView software (Bruker).

### Imaging data processing

#### Motion correction and ROI extraction

All raw imaging data were processed offline using Suite2p (available at https://github.com/MouseLand/suite2p, Pachitariu et al. 2016) for motion correction, region-of-interest (ROI) detection, and fluorescence trace extraction. For each imaging session: Motion correction: We first applied rigid and nonrigid registration to correct for x-y tissue motion. Typically, default Suite2p parameters were used.

ROI detection and classification: Suite2p’s built-in clustering algorithm was used to segment putative somatic ROIs based on pixel correlation and temporal fluorescence profiles. The algorithm automatically discarded obvious neuropil and blood vessels based on shape and time series criteria. For each candidate ROI, we visually inspected the ROI footprint and calcium trace to ensure it corresponded to a real neuron rather than a noise component. Typically, 50-150 ROIs were detected per plane, depending on GCaMP7f expression levels and imaging depth.

#### Trace extraction

For each accepted ROI, Suite2p extracted a raw fluorescence trace F(t) by averaging the pixel values within the ROI.

#### Identification of tdTomato-positive neurons

To label projection-specific neurons, we used a second imaging channel capturing tdTomato fluorescence. We ran Suite2p in a two-channel mode so it performed ROI segmentation on both GCaMP (green) and tdTomato (red) channels.

#### tdTomato ROI matching

For each GCaMP ROI, we computed the mean tdTomato fluorescence across the same pixels. We normalized these values within the recording session to identify outliers (e.g., z-score > 3). Any ROI with significantly elevated tdTomato signal was classified as a putative projection neuron.

#### Crosstalk removal

We calculated the Pearson correlation between each ROI’s GCaMP trace and its tdTomato trace across the entire session. If the correlation exceeded 0.5, we flagged that ROI for potential bleed-through or motion artifact. Those ROIs were removed from the tdTomato-positive pool unless manual inspection confirmed they were real dual-labeled neurons. This conservative threshold minimized false-positive labeling due to spectral overlap (**Figure S2**).

#### Population definitions

OB-p or mPFC-p neurons: GCaMP+ somata with robust tdTomato expression (resulting from retrograde AAV from OB or mPFC). Unlabeled (“untagged” or “unknown-projection”) neurons: GCaMP+ somata lacking tdTomato expression, hence the projections of which are *unknown*. “Any” neurons: refers to *any* GCaMP+ somata, including both labeled and unlabeled neurons. This is often used under random sampling as an unbiased representative of piriform cortical neurons.

#### Longitudinal tracking

For the GNG dataset, we utilized ROICaT (https://github.com/RichieHakim/ROICaT) on Suite2p segmentation results to track neurons across sessions (including the first passive days and three task days). Afterwards, for each animal, we further manually refined the longitudinal tracking results from ROICaT with a custom GUI in Python, with the objective to maximize correct tracking of labeled neurons. For analyses using longitudinal tracking, neurons were considered OB-p or mPFC-p neurons if they had tdTomato expression for at least 3 out of 4 days. This allowed us to analyze tracked neurons with at least 40 neurons per projection target (46 OB-p, 41 mPFC-p). See **Figure S10A** for illustration.

### Analysis

#### Fluorescence normalization

After extracting the raw fluorescence trace *F*[*t*] for each neuron, we calculated the normalized fluorescence trace by first computing the running baseline *F*_0_[*t*], which is estimated by first median-filtering *F*[*t*] over a 1-second window, then taking the 10th percentile over a 30-second moving window to capture slow baseline fluctuations. The normalized fluorescence is the difference between *F*[*t*] and *F*_0_[*t*], divided by the standard deviation of the difference, i.e., 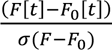.

#### Event detection (for 30Hz fast imaging)

In experiments with 30 Hz frame rate, we performed a peak-finding step to identify discrete calcium “events” for each trial, with the first and last seconds of all trials always ignored to minimize potential artifacts introduced by deconvolution. Specifically, for each neuron, calcium peaks were detected from normalized fluorescence traces with the following minimum criteria: 100ms calcium transient width, 1-second distance between peaks, normalized fluorescence height=0.5 and peak prominence=0.25.

Additionally, due to slow calcium dynamics, we also deconvolved using OASIS (79); AR(1) with L_0_ penalty and automated estimated *g*), then z-scored (clipping between −20 and 20 SD over mean). As a closer proxy to the onset of the calcium events, we detected the peaks of the z-scored deconvolved traces with the following criteria: 1-second distance between peaks, peak height=1 and peak prominence=2.

#### Projection decoding analysis with a transformer model

To build a robust neuron classification model, we considered two main challenges: (i) the limited number of available data, and (ii) the need to capture temporal patterns shared across neural recordings. Our approach addresses both by combining trial-level preprocessing, temporal patching, and transformer-based modeling. Supplement Figure S3A summarizes the overall pipeline, from time-series processing of a neuron to its final classification.

#### Preprocessing

The dataset consists of neural time-series recordings, each labeled with a corresponding class. We only used the *identity* passive dataset and the Go/NoGo dataset for the analysis involving the transformer. To ensure consistency, all trials were restricted to a fixed time window: 1 second before odor onset and 15 seconds after, resulting in 70 frames. Given the relatively small number of known projection neurons, we also augmented the dataset to enhance robustness and reduce overfitting: L=5 independent subsamples were constructed per neuron by randomly sampling M=20 trials from the original set of trials of that neuron.

#### Embedding

To efficiently model temporal structure and maintain robustness to small trial-level variations, each trial was divided into patches of p=2 consecutive frames. Each patch was then projected into a d-dimensional (d=128) latent space via a linear embedding, transforming the raw time-series into a sequence of patch embeddings, analogous to tokenized words in natural language processing. In this representation, each patch would serve as a token encoding localized temporal information.

#### Model

We adopted a Vision Transformer (ViT)-based architecture (80) for temporal modeling. The sequence of patch embeddings is treated as input tokens, with a learnable class token appended at the beginning of each sequence. This token aggregates global information across all patches and ultimately is used for classification. Within the transformer, multi-head self-attention captures temporal dependencies across patches.

#### Interpretability

Importantly, by analyzing attention weights associated with the class token with attention rollout (42), we could identify which temporal segments contribute most strongly to the classification decision. This provides interpretability, highlighting temporal dynamics relevant for distinguishing neuron classes.

#### Training

After transformer processing, we obtained latent representations for the M=20 trials within each of the L=5 subsamples. These trial-level embeddings were average-pooled to produce a stable neuron-level representation. Since multiple subsamples per neuron were generated, their predictions were aggregated using majority voting, yielding the final class label for each neuron. The transformer was trained using the hyperparameters reported in **Table 5**. We used the Adam optimizer (81) with a cosine decay scheduler, which periodically decreases the learning rate following a cosine function.

**Table 5:** Hyperparameter Settings for Transformer Model.

| Hyperparameters | Value |
| --- | --- |
| Batch Size | 8 |
| Learning Rate | 1e-3 |
| Epochs | 200 |
| Number of Samples (L) | 5 |
| Trials per Sample (M) | 20 |
| Timeframes per Patch (p) | 2 |
| Embedding Dimension (d-dim) | 128 |
| Transformer Depth | 6 |
| Attention Heads | 4 |
| MLP Expansion Factor | 2 |
| Transformer Dropout | 0.2 |

#### Evaluation

We evaluated performance in two settings:

- Within-animal evaluation: Neurons from all animals were pooled together and randomly split into training/validation/test sets with ratios 0.8/0.1/0.1. This setting measures the model’s general classification performance across neurons.
- Across-animal evaluation: To assess generalization across animals, we held out pairs of animals for testing. Each pair was drawn from the same dataset (Identity or GNG), with one animal having OB projections and the other mPFC projections. Pairs were chosen to ensure approximately balanced neuron sizes between animals (**Table 6**).

All results were averaged across multiple random seeds: 10 seeds (0, 1, 2, 3, 4, 42, 43, 44, 45, 46) for within-animal evaluation, and 3 seeds (42, 43, 44) for across-animal evaluation. The seeds were applied consistently to both the model initialization, the data splits, and random generation.

**Table 6:** Animal Pairs Selected for Across Animal Evaluation.

| Dataset | OB-p tagged subjects (# neurons) | mPFC-p tagged subjects (# neurons) |
| --- | --- | --- |
| Identity | 8 (10) | 533 (8) |
|  | 9 (12) | 433 (14) |
|  | 164 (15) | 512 (14) |
|  | 163 (22) | 448 (17) |
| GNG | 664 (96) | 2456 (107) |
|  | 2400 (134) | 2457 (131) |

#### Alternative model (control)

To verify that the transformer’s performance comes from its ability to capture temporal dependencies via self-attention, we conducted a control experiment by replacing the transformer with a multilayer perceptron (MLP).

We performed a Bayesian hyperparameter sweep over 50 runs using Weights & Biases, with the hyperparameter search space reported in **Table 7**. The best MLP configuration was selected based on validation performance in the within-animal setting (seed 42). The final chosen hyperparameters are reported in **Table 8**.

**Table 7:**
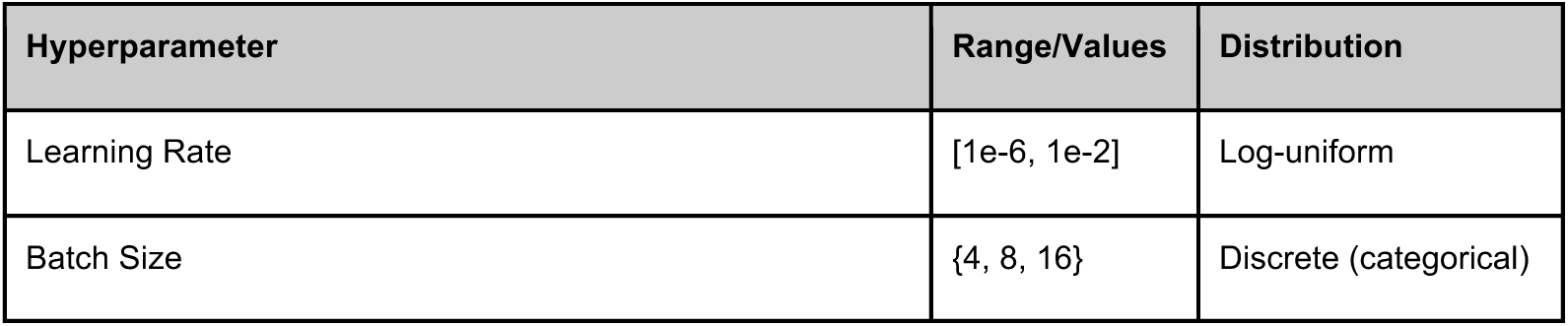
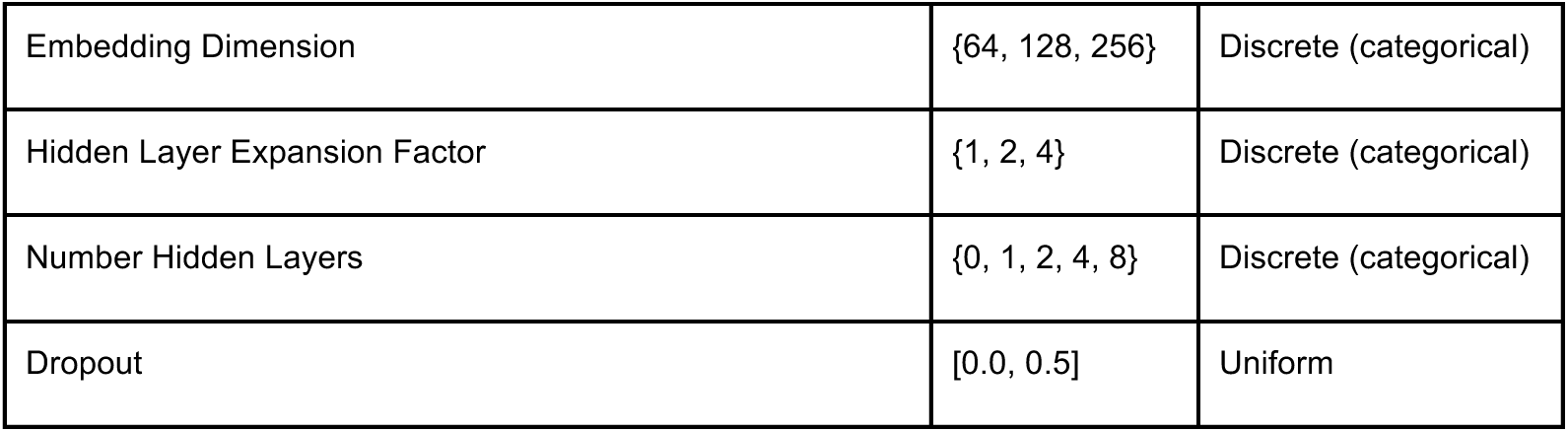
Hyperparameter search space used in the MLP sweep experiments.

| Hyperparameter | Range/Values | Distribution |
| --- | --- | --- |
| Learning Rate | [1e-6, 1e-2] | Log-uniform |
| Batch Size | {4, 8, 16} | Discrete (categorical) |
| Embedding Dimension | {64, 128, 256} | Discrete (categorical) |
| Hidden Layer Expansion Factor | {1, 2, 4} | Discrete (categorical) |
| Number Hidden Layers | {0, 1, 2, 4, 8} | Discrete (categorical) |
| Dropout | [0.0, 0.5] | Uniform |

**Table 8:** Final Hyperparameters for MLP experiments.

| Hyperparameter | Value |
| --- | --- |
| Learning Rate | 5.85e-4 |
| Batch Size | 4 |
| Embedding Dimension | 128 |
| Hidden Layer Expansion Factor | 4 |
| Number Hidden Layers | 1 |
| Dropout | 0.264 |

#### Determination of odor-responsive (“active”) neurons

For certain analyses (e.g., cellular properties), we established a binary indicator of response presence or absence on each trial: Baseline Distribution: To accommodate non-parametric statistical tests for active response determination, we first sampled 1000 points of the normalized trace within 4 seconds before odor onset across all trials to build a null distribution of baseline values (X*_b_*).

#### Response Window Criterion

We then examined each trial’s trace from the first 5 seconds (0–5) after odor onset (or the first 2 seconds (0–2) for certain analyses; in GNG dataset, we performed this binarization step for each task-relevant phase). A forward sliding one-sided Mann-Whitney U test (width = 0.7 s) was performed, comparing the response window of interest to X*_b_*. If p < 0.01 for at least 8 consecutive forward frames, the neuron was considered “forward-active” in that trial. This approach follows Wang et al. (2020) for robust detection of odor-evoked responses in noisy calcium signals.

#### Pre-odor Exclusion Criterion

For response windows that start at odor presentation (e.g. first 5 seconds in Identity dataset or odor phase in GNG dataset, but not delay phase in GNG dataset), we also performed similar one-sided Mann-Whitney U tests against the baseline distribution, but in a backward sliding manner. If p < 0.01 for any of the 4 consecutive backward frames, the neuron was considered “backward-active”.

#### Finalized Active Criterion

For a given response window that starts after odor onset (e.g. act-early phase in GNG dataset), the neuron was considered “active” in a trial if it was already determined “forward-active” in that trial. For a response window that starts at odor onset (e.g. first 5 seconds), the neuron was considered “active” in a trial only if it was both “forward-active” and *not* “backward-active”. The exclusion criterion was applied to avoid potential false positives of left-over spontaneous calcium transients preceding odor presentation.

As a result, for each neuron *n*, if it is active in trial *k*, we denote the binary activity as 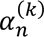 = 1. Otherwise, we denote as 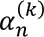 = 0.

#### Metrics derived from response binarization

*Mean (binarized) activity* of a given cell *n* is the proportion of trials that the cell is active, i.e., ∑ 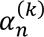/*K* where *K* is the total number of trials.

*Proportion of active cells* for a given trial *k* and a given population of neurons {*n_i_*,} (e.g. only OB-p neurons) is defined as the proportion of active cells for that trial, i.e. ∑ 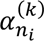⁄*N* where *N* is the number of neurons in that population.

*Lifetime sparseness* of a given cell *n* describes the tuning width of the neuron to the different stimuli {*s_j_*}. For each trial *k*, we have a stimulus *s*^(*k*)^ ∈ *S* = {*s_j_*} and *k_s_* = |*S*| as the number of stimuli. Define 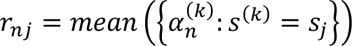 as the trial-averaged activity of the cell *n* for each stimulus *s_j_*. Then the lifetime sparseness is defined as:

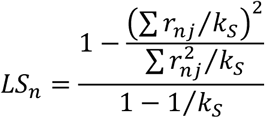

*Population sparseness* of a given population of neurons {*n_i_*} (and the total number of cells is *N*) for a given stimulus *s* can be calculated similarly. Define 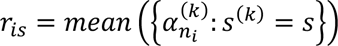 as the trial-averaged activity to the stimulus *s* from each cell *n*,. The population sparseness is defined as:

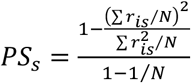

*Mutual information* between a neuron’s binarized activity *n* and a stimulus set *S* describes the nonlinear mutual dependence between the neuron’s binarized activity and the stimulus distribution. Denote *H*(*X*) as the entropy of the discrete variable *X*.

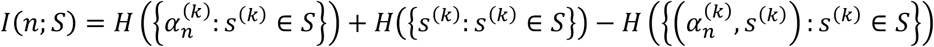

#### Reliability analysis

We used the same procedures as in our earlier work to classify neurons as reliable or unreliable (82). Briefly, for each cell, we calculated the mean baseline activity before odor onset and identified a neuron as responsive if the mean activity during a defined window after odor onset (0–5 s) exceeded a threshold. This threshold was determined by adding a set number of baseline standard deviations (SDs) to the mean baseline. The number of SDs was chosen according to the significance level, which in this case was p = 0.01, corresponding to 2.33 SDs above baseline.

Reliability was assessed on a trial-by-trial basis in two steps. First, for each cell, we examined all trials and classified the cell as reliable if it responded in more than half of them, and unreliable otherwise. In a second step, we quantified for each trial the number of reliable and unreliable cells. These values were used in the reliability analyses and figures.

#### Pseudo-population construction

For each cell and trial, given a certain time window, we aggregated the response by using either the average or maximum of the normalized fluorescence trace. For each projection population (OB, mPFC, unknown/untagged), we constructed pseudo-population response vectors by combining cells across subjects. All bad trials (e.g. trials with extreme fluorescence values due to the microscope not recording) and control odor trials (e.g. stimuli such as mineral oil or air) were removed beforehand. The trials were then concatenated across animals, such that each resulting pseudo-population trial would (a) come from trials with the same stimulus identity and (b) have the same trial order of the stimulus as the originating trials from each subject, i.e. the first trial of each stimulus would be combined across subjects, and not necessarily the first trial of the session from each subject. Additionally, as the total number of labeled neurons per projection target varies, we also subsampled from the pseudo-population randomly without replacement to compare equally sized pseudo-populations from different projection targets. For the Go/NoGo dataset, we focused our analyses on task days, hence pseudo-populations would be constructed for each task day separately.

#### Population correlation

Pseudo-population responses (i.e., a trial-by-cell matrix) are used to calculate population Pearson correlation across trial pairs (i.e., a trial-by-trial matrix). Stimuli-pair correlations are computed by averaging the relevant blocks in the trial-by-trial correlation matrix. All correlations within the exact same trial are discarded before averaging the population correlations within similar-stimulus trials, because they would always be exactly 1.0 and would create bias in the correlation average.

#### Decoding analysis

For all decoding tasks, we used a linear SVM with all default parameters as given by *sklearn*.

For the *identity* dataset, we first trained and tested the classifier to decode the 8 odor identities using pseudo-population responses from mean-aggregate between 1 and 5 seconds after stimulus onset, with 5-fold cross-validation. We repeated this process with varying subsampling sizes. To investigate potential decoding dynamics, we also trained and tested the classifier on various pairs of time windows, using maximum 75 cells to subsample from, each of which is 500ms long. To avoid data contamination, especially because of calcium dynamics, train and test samples were never on the same trial, even if the train and test time windows were different.

For the *concentration* dataset, we also first train the classifier to decode the 9 unique odor (i.e. combination between odor identities and concentrations) pseudo-population responses from mean-aggregated between 0 and 5 seconds after stimulus onset, with 5-fold cross-validation. Variation of pseudo-population size was also done for this choice of window. To investigate decoding dynamics, we also train/test on different time windows, as previously. To investigate specifically concentration decoding dynamics, for each identity class (i.e. odorant), we train a classifier to decode concentration levels, and test on different time windows; then we average the decoding accuracy matrices across different stimulus classes for visualization.

For the *Go/NoGo* dataset, there were six separate classifiers:

1. odor identity: Decode the 4 conditioned stimuli (CS) from all trials.
2. odor identity for correct trials: Decode the 4 CS on only correct trials (for both training and testing).
3. valence (unconstrained): Decode the stimulus valence (i.e. *reward* for CS*k*1 and CS*k*2 and *none* for CS-1 and CS-2) from all trials, and train/test splitting is *not* dependent on odor identity sources.
4. generalized valence: Decode stimulus valence by splitting by odor identity sources, e.g. train on CS*k*1, CS-1, then test on CS*k*2, CS-2. There are in total 4 permutations of this train/test splitting paradigm.
5. valence (unconstrained) for correct trials: Decode stimulus valence for *correct* trials.
6. generalized valence for correct trials.

With pseudo-populations in the Go/NoGo dataset, we performed decoding analysis for each session separately. Within each session, we evaluated using either discrete non-overlapping running 1-second time windows or task-relevant phases (see below). Pseudo-population analyses were performed with either train/test on the same window (or phase) or train/test across different combinations of discrete windows (or phases).

For each decoding analysis, a pseudo-population was constructed by randomly sampling 85 cells for 100 times for each projection target: OB-p, mPFC-p, or any-p. Details about decoding analysis with tracked projection neurons are in the later section.

#### Task-relevant phases of GNG dataset

For each trial in the GNG dataset, analyses were split into 4 task phases (see **Figure 4B**):

- odor (abbreviated as O in **Figure 5**): 0-2 sec after stimulus onset
- delay (abbreviated as D in **Figure 5**): 2-5 sec after stimulus onset
- act-early (abbreviated as AE in **Figure 5**): 5-10 sec after stimulus onset
- act-late (abbreviated as AL in **Figure 5**): 10-15 sec after stimulus onset

#### Session-to-session analyses of metrics

The session-to-session analyses of single-cell metrics are only relevant for the persistently tracked labeled neurons across task days (Days 1-3). For each session and each task phase, in addition to the aforementioned single-cell metrics derived from binarized activity, we also quantified the averaged response as the mean of the fluorescence traces across trials, and the trial-to-trial variability as variance of the fluorescence across trials. To capture the effects of learning, for each metric across sessions, we quantified the slope and the coefficient of variation of the metric. While the former is informative of the monotonic direction of change across learning, the latter is informative of the plasticity of the cells across learning.

#### Decoding with tracked neurons across Go/NoGo sessions

For each subject, we performed decoding analyses from activity of ensembles of persistently tracked and labeled neurons. Specifically, we built various classifiers, as laid out above, to quantify variations of identity and valence information capacity. To assess how transferable ensemble information is across learning, we performed training and testing on different combinations of pairs of tasks phases and task days. These form transferability heatmaps for each subject, which we also averaged across animals.

Then, we quantified how decoding accuracy changed when trained and tested within the same sessions, across proximal (e.g., Day 1 and Day 2) or distal (Day 1 and Day 3), in which only adjacent phases (e.g., delay and act-early) would be considered to avoid grouping analysis of discontinuous phases (e.g., odor and act-late) (**Figure 5F**; additionally see **Figure S12** for other considerations of phase or day difference). This would inform how far information could be transferred across learning (see **Figure S11B** for demonstration).

Finally, we quantified the stability of temporal structure of these information contents by computing the pairwise correlations of session-to-session task-phase transfer blocks (i.e. each internal 4-by-4 matrix in **Figure 5E**). Higher Pearson correlation coefficients would entail higher stability (see **Figure S11B** for demonstration and difference between transferability and stability concepts used and computed in this study). See also **Figure S13** for usage of other correlation methods.

### Statistics

Details of statistical tests and adjustments are stated in the text. For multiple comparison correction, we used Bonferroni correction for the majority of analyses, with the exception of comparisons concerning single-cell properties of tracked neurons, which we used Benjamini-Hochberg due to the lower number of cells to be less conservative during exploratory analysis concerning tracked neurons.

---

**Figure S1.**
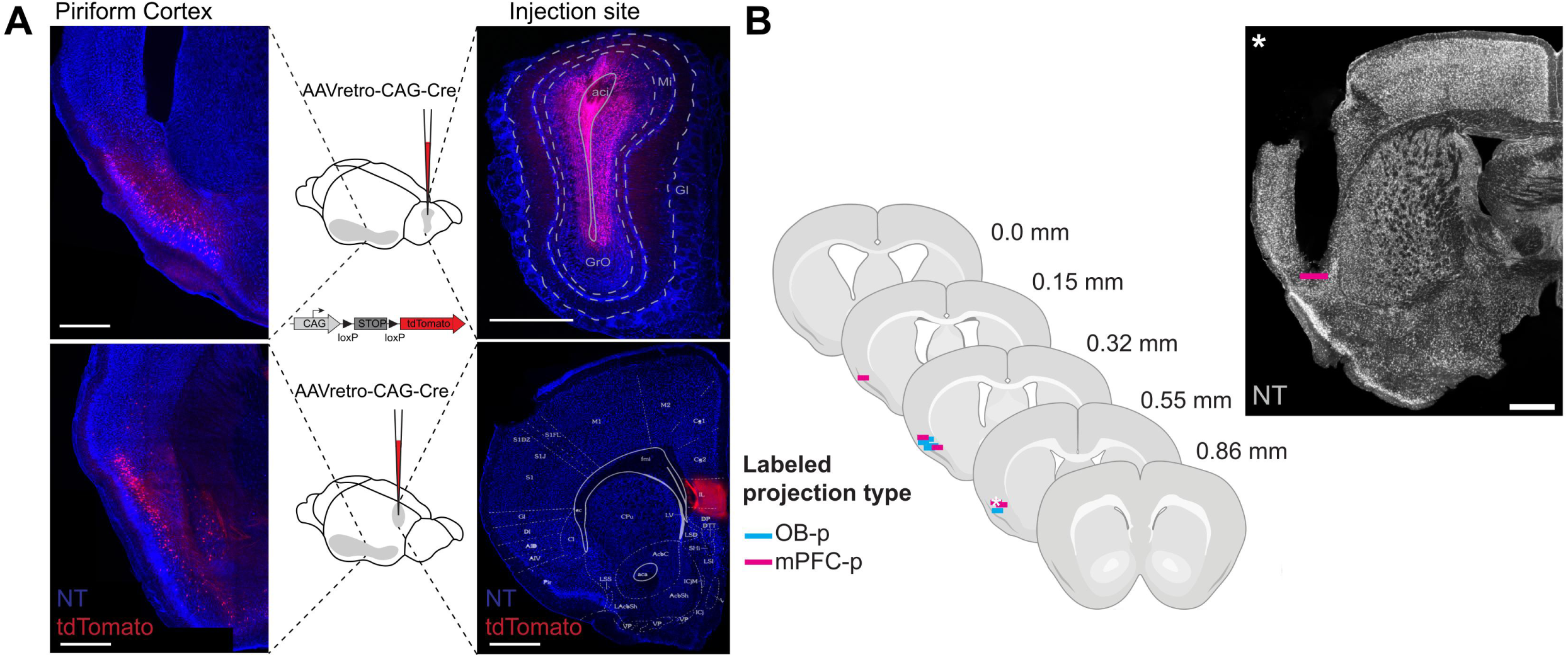
Histological characterization of the animals used for imaging. **(A)** Injection of AAVretro-CAG-Cre only in Ai14 mice with injection represented on the right and resulting labeling in PCx on the left, either in OB (top) or mPFC (bottom). Scale bars: 500μm. (**B**) Histological confirmation of GRIN lens placement centered around 0.3 mm on the anterior-posterior axis in both experimental mouse conditions (5 OB-p tagged mice; 5 mPFC-p tagged mice; represented mice are from the identity dataset). Star shows placement of one mouse with a related coronal section on the right. Mice injected with AAVretro-CAG-Cre in OB and mPFC are represented in cyan and magenta, respectively. Scale bar: 500μm.

**Figure S2.**
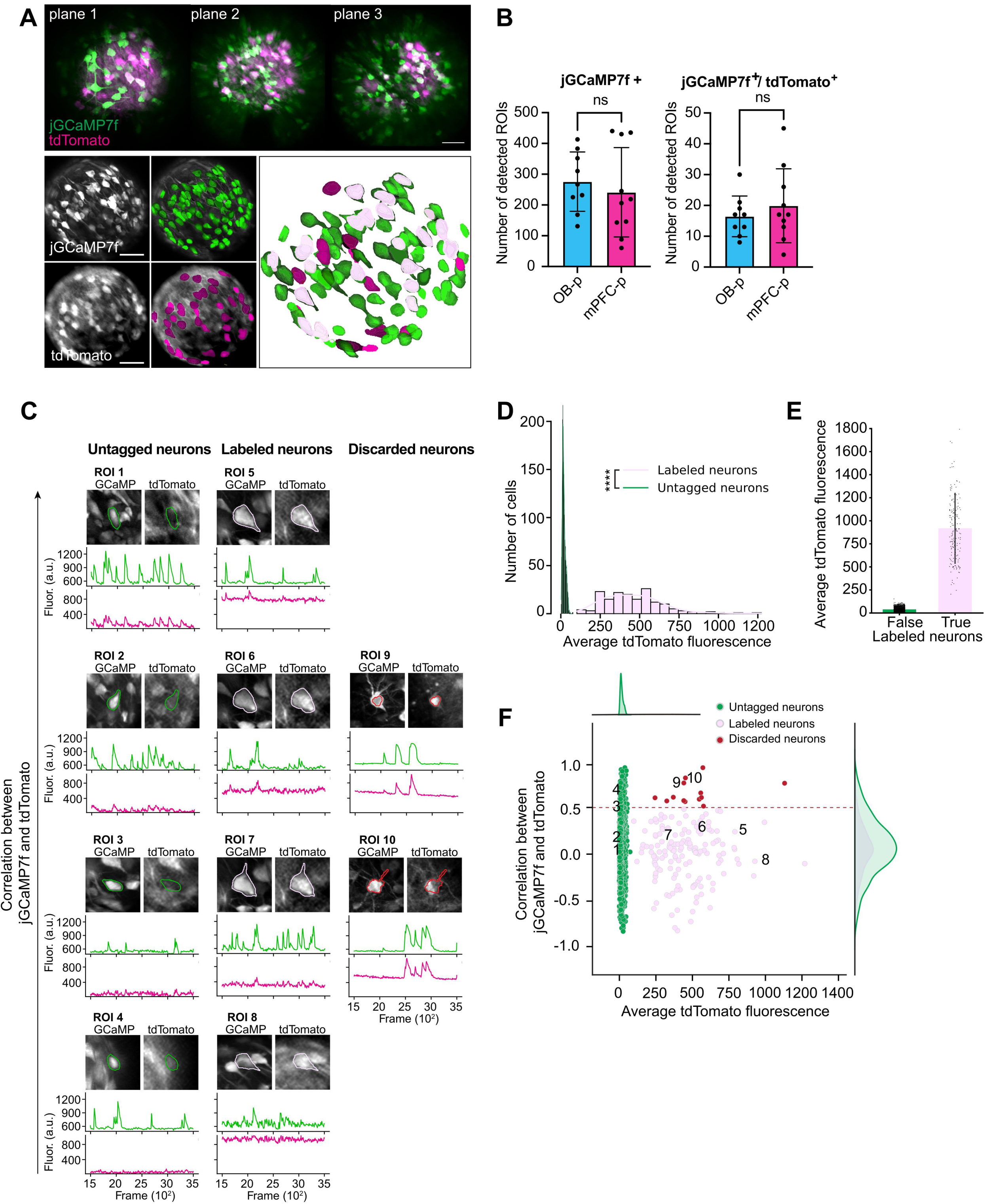
PCx projection neurons co-expressing jGCAMP7f and tdTomato are identified *in vivo*. (**A**) Top: Maximum projection of the field of view during in vivo imaging through the GRIN lens. jGCaMP7f (green) and tdTomato (magenta) are simultaneously imaged in 3 planes on the z-axis. Scale bar: 200 μm. Bottom: Example of segmented ROIs. Bottom left: maximum intensity z-projections of jGCaMP7f and tdTomato signal. Scale bar: 200 μm. Middle: Overlay with segmented ROIs for jGCaMP7f and tdTomato. Right: Overlay of jGCaMP7f positive and tdTomato positive ROIs, with double positive ROIs in light pink corresponding to labeled projection neurons. (**B**) Quantification of jGCaMP7f and co-labeled jGCaMP7f/tdTomato-expressing neurons in OB-p and mPFC-p mice. (OB-p, N = 9; mPFC-p, N = 10. Unpaired two-tailed t-test: left: p=0.555; right: p=0.455). (**C**) Representative examples of untagged, labeled, and discarded neurons based on the correlation between jGCaMP7f and tdTomato fluorescence. For each neuron, the top panels show the segmented ROI overlaid on the jGCaMP7f (green) and tdTomato (magenta) mean images, and the lower panels display the corresponding fluorescence traces across time. Untagged neurons (left) exhibit strong jGCaMP7f activity and minimal tdTomato fluorescence. Labeled neurons (middle) show concurrent jGCaMP7f and tdTomato signals consistent with projection neuron identity. Discarded neurons (right) display high jGCaMP7f–tdTomato correlation (r > 0.5), indicating false-positive co-labeling. (**D**) Distribution of average tdTomato fluorescence signal in all ROIs, normalized by the distance from the center of the imaging FOV (p-value: 3.10E-68, unpaired t-test).(**E**) Comparison of average tdTomato fluorescence for labeled and untagged neurons. (**F**) Correlation between jGCaMP7f and tdTomato fluorescence plotted against average tdTomato signal. Each point represents a single neuron, color-coded as in (**C**). The dashed red line indicates the correlation threshold (r = 0.5) used to exclude false-positive co-labeled neurons. Numbers indicate the example ROIs shown in (**C**).

**Figure S3.**
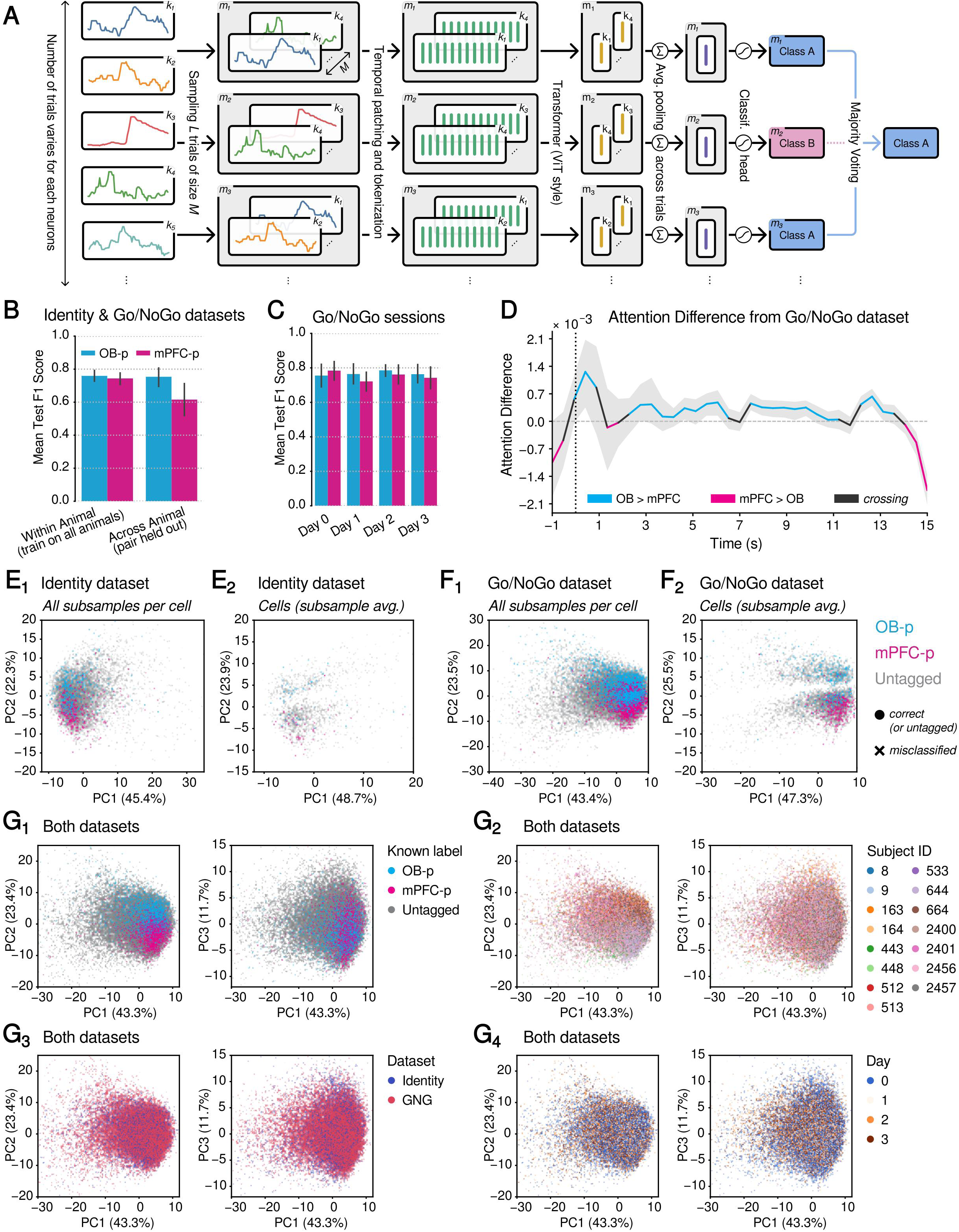
**Extended results and analyses of the transformer used to classify PCx projection targets based on neural activity**. (**A**) Detailed schematic of the transformer-based classifier for predicting neuron projection targets from single-cell calcium dynamics. For each neuron, calcium traces were sampled across trials (*k_i_*) to form subsamples (*m_j_*). Within each subsample, trials were divided into non-overlapping temporal patches and tokenized for input to a transformer (ViT-style). Trial-level latents were averaged, pooled, and passed through a classification head to predict projection identity. Final neuron-level classification was obtained by majority voting across subsamples. (**B**) Evaluation of the transformer’s generalization capacity when training and testing were performed with data from *separate* animals (right bars), compared to the scenarios where train/test sets contained all animals (left bars). The transformer was trained and tested on the combined passive identity and Go/NoGo datasets (see **Methods**), for either 10 different seeds (left bars, “all animals” case) or 3 seeds per held-out pair (right bars, 3 seeds x 6 pairs = 24 instances). F1 scores were shown separately for OB-p labels (cyan) and mPFC-p labels (magenta). (**C**) Classification performance of the transformer trained on both datasets containing all animals and tested on different Go/NoGo sessions. The model was trained for 10 different seeds. (**D**) Attention-rollout analysis of the transformer model trained on both datasets and evaluated for the Go/NoGo dataset. Similar to Figure 1G, which was evaluated for the passive identity dataset. (**E**) Principal component analysis (PCA) of latent representations extracted by the transformer model first trained on both datasets then evaluated only on the passive identity dataset. Only the model from one seed (seed=42) was shown for demonstration. Each point in **E_1_** corresponds to the latent embedding of a single subsample (*m_j_*) after trial-averaged pooling. In **E_2_**, each point represents the latent embedding of a single neuron, obtained by averaging the latent representations of subsamples from **E_1_** that agree with the neuron’s majority classification outcome. In other words, subsamples that do not match the majority classification were excluded before averaging the previously trial-averaged latent representations. Note that in both instances (**E_1_** and **E_2_**), the first 2 PCs already capture around 70% explained variance. Points are colored by projection targets (cyan: OB-p, magenta: mPFC-p, grey: untagged) and classification outcomes (circle: correct, cross: incorrect; for known projection targets). (**F**) Same as **E**, but evaluated only on the Go/NoGo datasets. Similar to **E**, both instances (**F_1_** and **F_2_**), the first 2 PCs already capture around 70% of explained variance. Note that the PCA space for **E** and **F** were constructed using only the data from each of their respective dataset. For the PCA space constructed from the combination of both of these datasets, see **G**. (**G**) PCA of the latent representations (of each subsample) extracted by the transformer model first trained on both datasets then evaluated on both datasets. Three PCs are shown in these panels, capturing near to 80% of explained variance. The points are colored by different sources of metadata. **G_1_**: known projection labels. **G_2_**: subject IDs. **G_3_**: dataset sources. **G_4_**: Go/NoGo sessions (day=0 includes both the identity passive dataset and the passive session of the Go/NoGo dataset).

**Figure S4.**
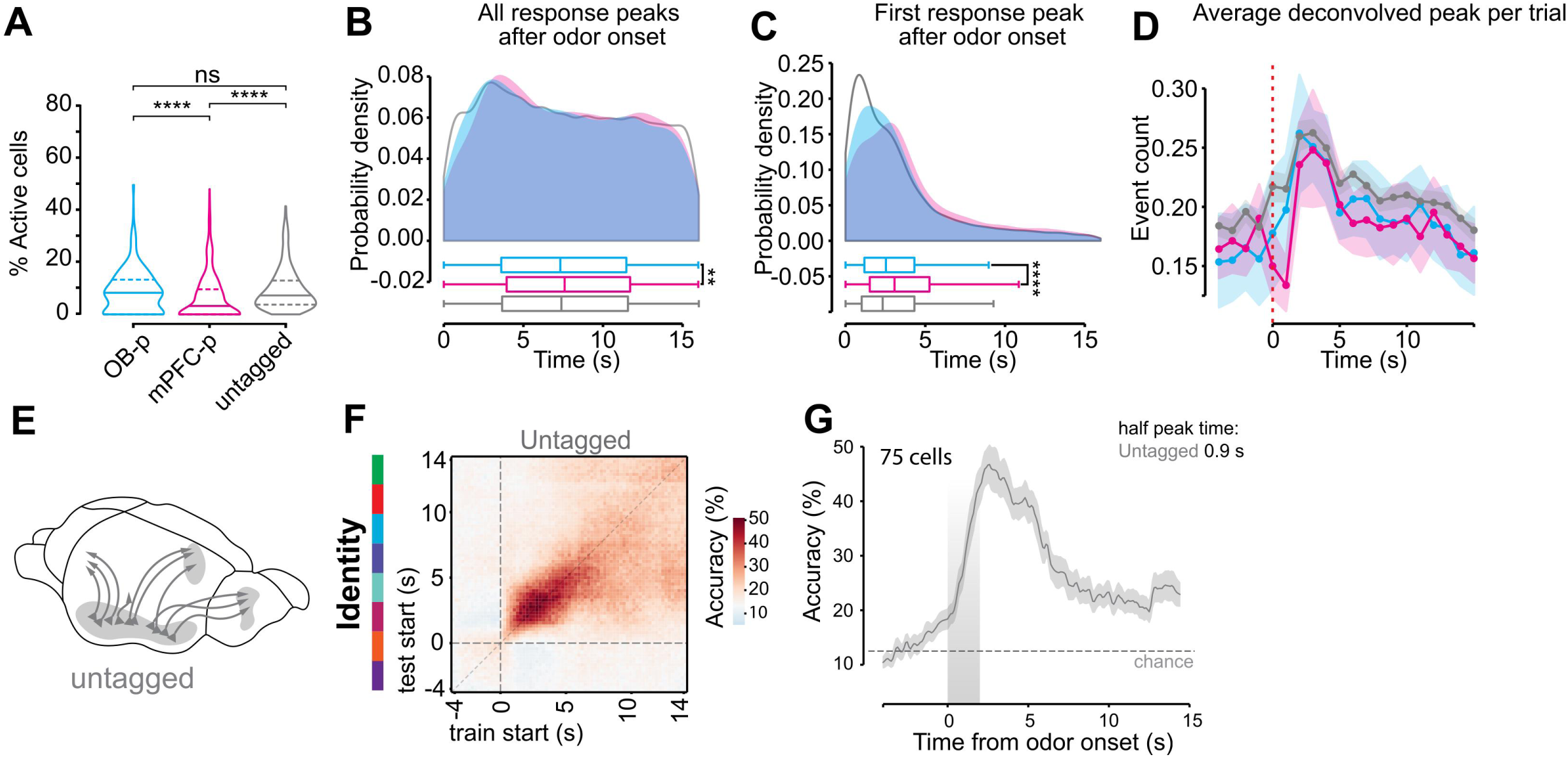
Extended odor response properties of OB- and mPFC-projecting PCx neurons from the fast (30 Hz) dataset. (**A-C**) These events are the peaks of the deconvolved traces using OASIS (see **Methods**), which can be considered as a closer proxy to the onset of calcium traces. Refer to Figure 2F and G for analysis of calcium peaks. (**D**) Schematic illustration of the untagged population of piriform cortex neurons used for comparison with projection-defined OB-p and mPFC-p neurons. (**E**) Cross-temporal decoding matrix for odor identity using untagged neurons. Linear SVM classifiers were trained on single-trial response vectors in 250-ms windows (x-axis) and tested on all windows (y-axis), using pseudo-populations of 75 cells (100 repeats). Color indicates classification accuracy. White denotes chance level (12.5%). (**F**) Diagonal decoding accuracy from (E), showing odor identity classification performance when training and testing on the same time window. Curve shows mean accuracy across pseudo-populations, with shading indicating 95% confidence intervals. The dashed line marks chance level. The half-peak decoding time for untagged neurons is indicated above the curve.

**Figure S5.**
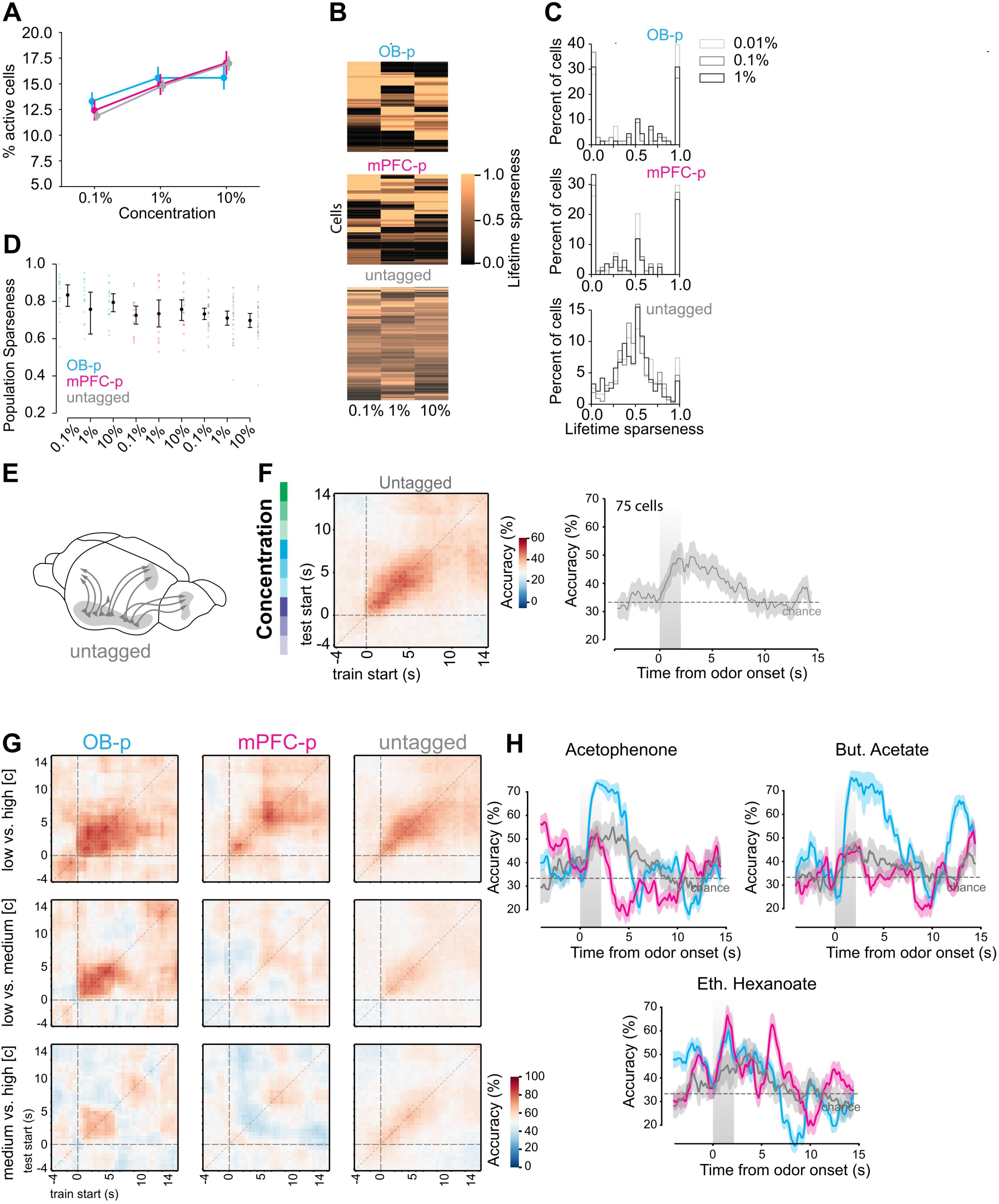
Additional analyses of concentration responses in OB-p, mPFC-p, and untagged neurons. (**A**) Percentage of active cells (fraction of trials with significant odor-evoked responses) for each projection type across three odor concentrations (0.1%, 1%, 10%). (**B**) Heatmaps of lifetime sparseness values for individual neurons at each concentration (0.1%, 1%, 10%) for OB-p, mPFC-p, and untagged populations. Each row is a cell; darker colors indicate higher sparseness. (**C**) Distributions of lifetime sparseness values for OB-p (top), mPFC-p (middle), and untagged neurons (bottom), shown separately for each concentration. (**D**) Population sparseness values for OB-p, mPFC-p, and untagged neurons across the three odor concentrations. Points show subject-level means ± SEM. (**E**) Schematic illustration of the untagged population used for comparison with projection-defined neurons. (**F**) Cross-temporal decoding of odor concentration using untagged neurons. Left: linear SVM classifiers trained and tested on 500-ms windows (x-axis = train time, y-axis = test time), using pseudo-populations of 75 cells (100 repeats). Right: diagonal decoding accuracy from the same analysis, showing mean ± 95% CI. Dashed line marks chance level (33.3%). (**G**) Cross-temporal pairwise concentration decoding (low vs. medium, low vs. high, medium vs. high) for OB-p (left), mPFC-p (middle), and untagged neurons (right). Heatmaps show accuracy across training (x-axis) and testing (y-axis) windows. Chance level is 50%. (**H**) Odor-by-odor concentration decoding accuracy for acetophenone, butyl acetate, and ethyl hexanoate. Linear SVM classifiers trained and tested on 250-ms windows using 75-cell pseudo-populations (100 repeats). Curves show mean accuracy ± 95% CI; dashed lines denote chance level.

**Figure S6.**
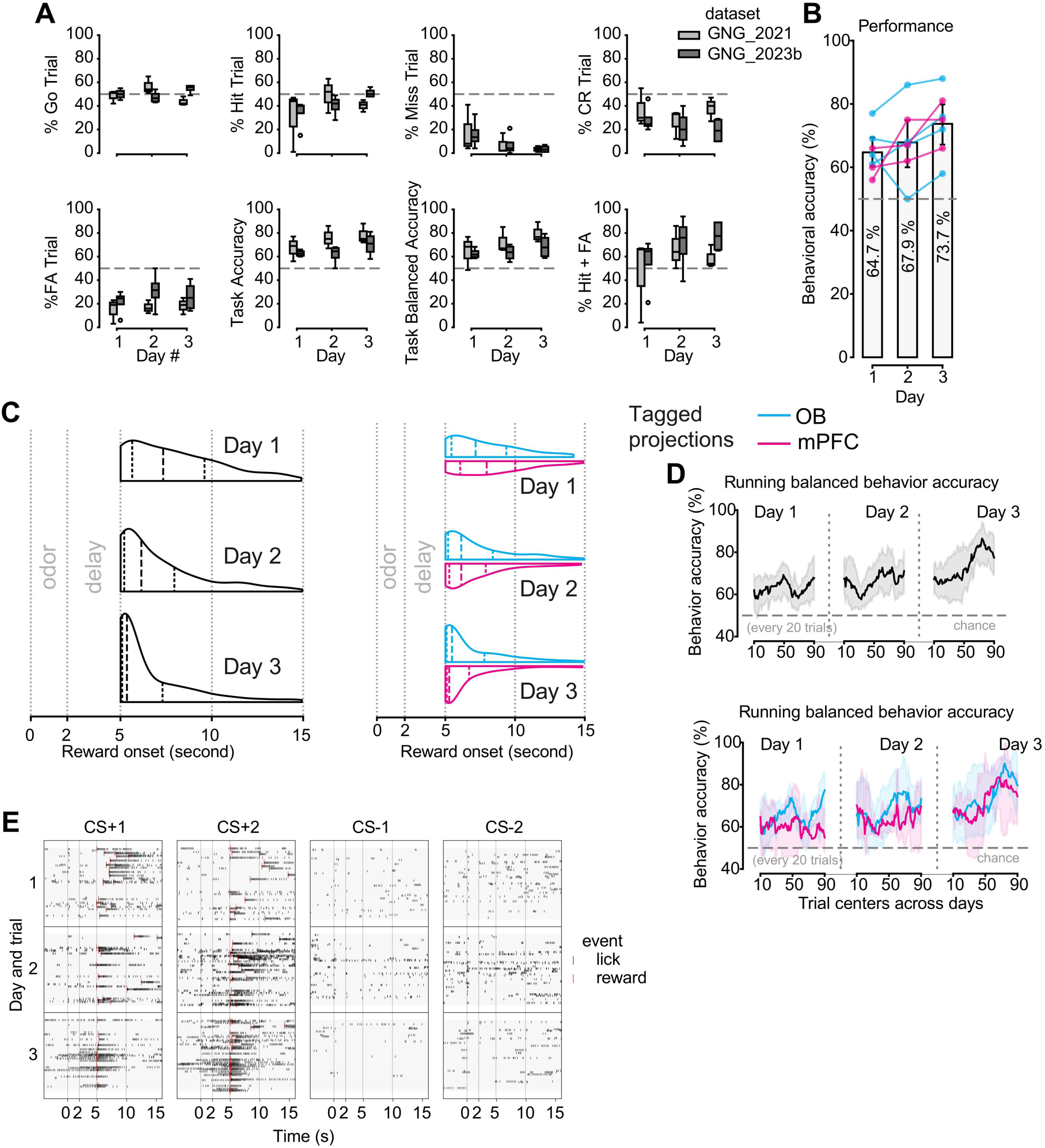
Extended Go/NoGo behavioral analyses. (**A**) Percent trials and performance across days of different Go/NoGo datasets. (**B**) Behavioral accuracy plotted as in Figure 4C (left), with additional information about animal batches. (**C**) Reward onset (Hit trials) compared across days and tagged projections. Note that this is a sufficient proxy for lick latency after delay for rewarded licks. (**D**) Balanced behavior accuracy ((TPR *k* TNR) / 2) with running trials (20-trial window) across days. (**E**) Example lick behaviors across learning. Rows (y-axis) represent trials and days; columns separate conditioned stimuli. Black ticks: lick events; red ticks: reward delivery.

**Figure S7.**
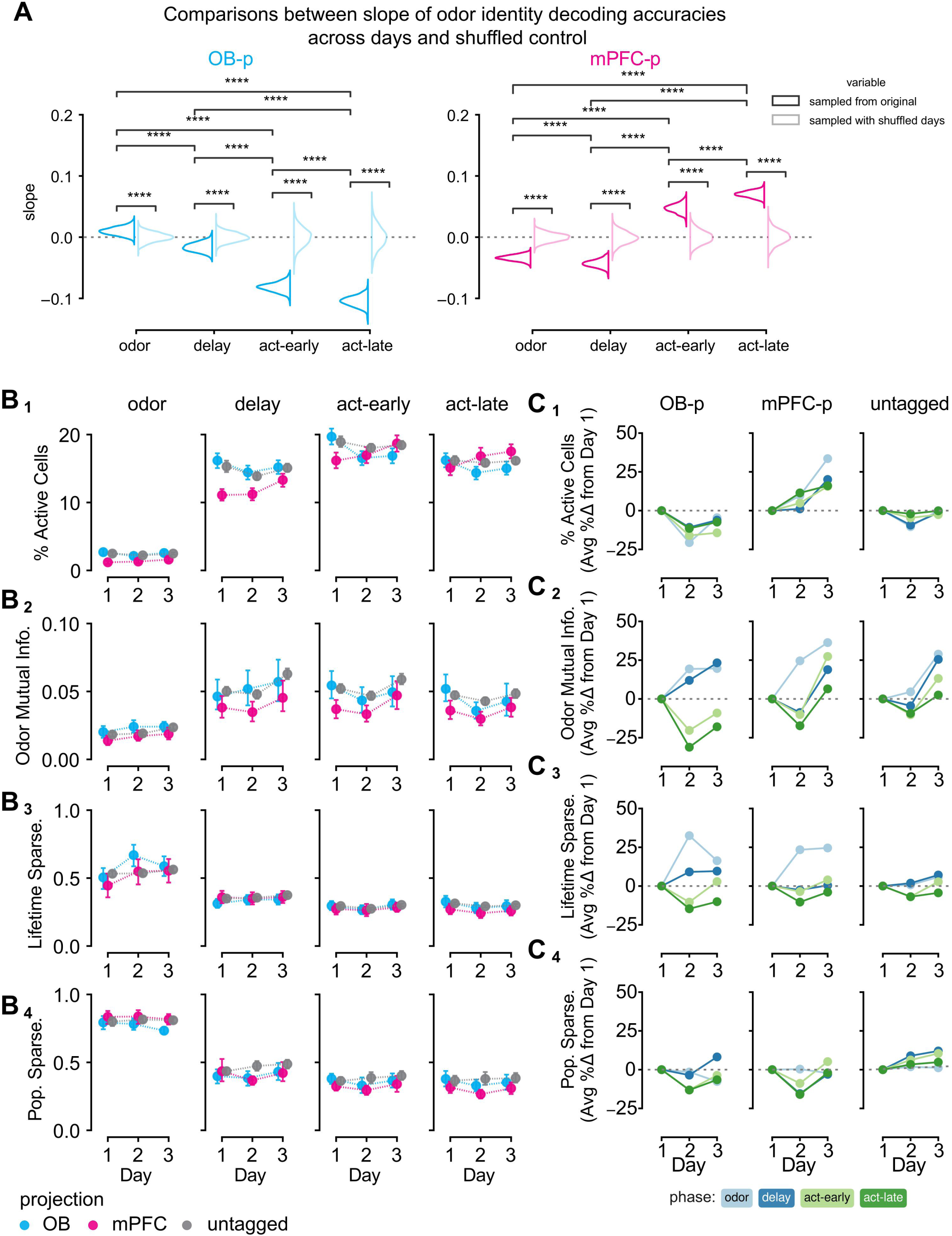
Learning effects on cellular properties and identity decoding. (**A**) Each slope is the slope of accuracy over days, in which the identity decoding accuracies were randomly sampled from 50 size-matched pseudo-populations for each task phase (x-axis) and projection type (panels). Days were also shuffled to establish null distribution (grey), to be compared with slopes sampled from the original accuracies (black). Each distribution was constructed by sampling 1000 times. Two-sided Mann-Whitney U tests with Bonferroni corrections were performed. (**B, C**) Cellular and population properties (top to bottom) quantified for different task phases (left to right panels in **B**, colors in **C**) across days (x-axis) of different cell groups (colors in **B**, left to right panels in **C**). Plots in (**B**) show the quantified metrics and those in (**C**) show the normalized metrics (first averaged in (**B**) then normalized to obtain the % difference relative to Day 1).

**Figure S8.**
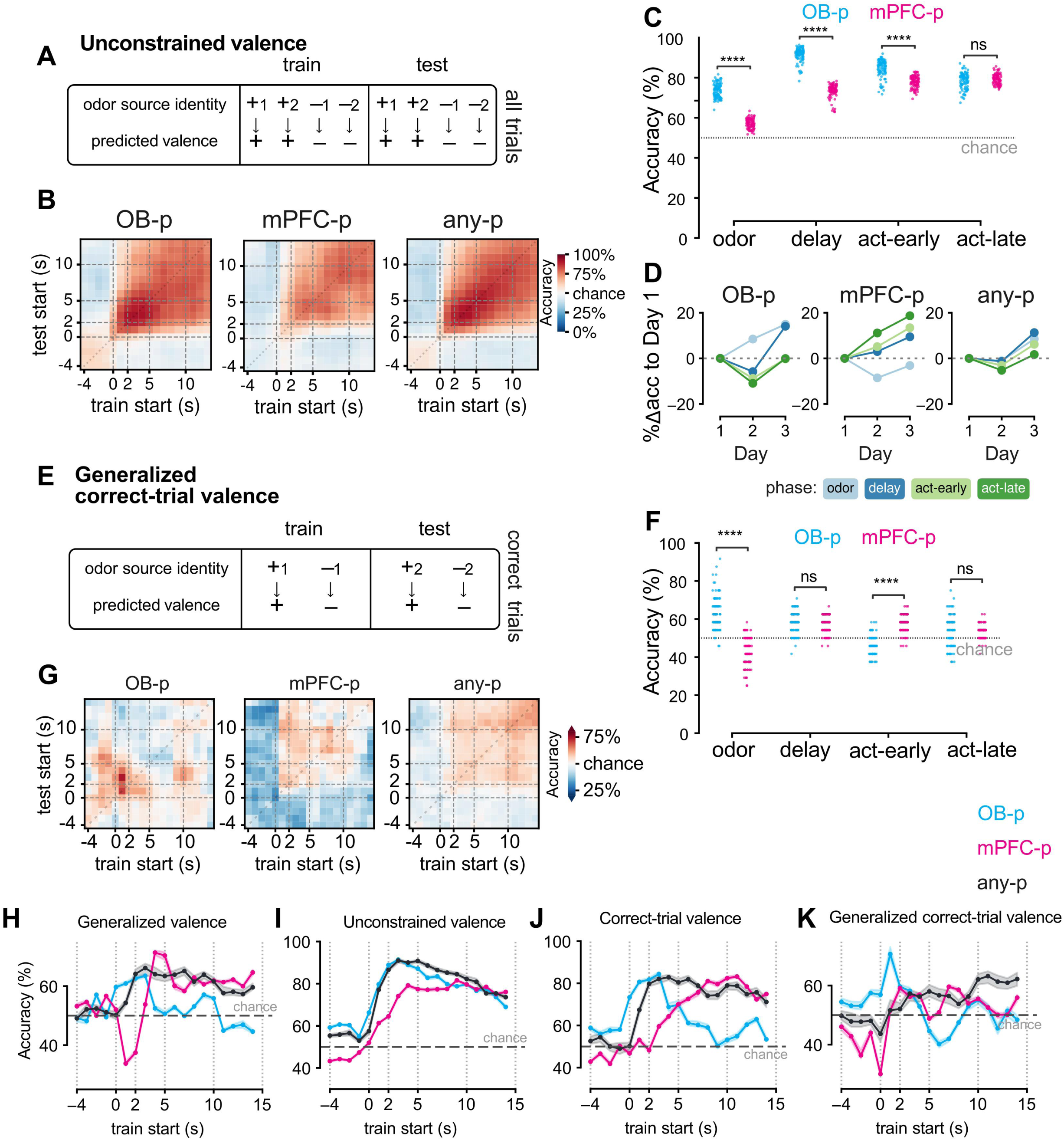
Extended valence decoding analysis for Day 3 of Go/NoGo task. (**A-C**) Unconstrained valence decoding in Day 3, and across learning (**D**), of the Go/NoGo task. See Figure 4G and H for a similar description. The difference between generalized and unconstrained valence is that the former means the model must learn the higher-order categorization of odors by training/testing on trials of different odors, whereas the latter does not require odor identities to perform train/test splitting. (**E-G**) Generalized valence encoding for correct trials in Day 3 of Go/NoGo task. (**H-K**) Valence decoding accuracy when pseudo-populations were trained and tested on the same 1-second time window within a trial in Day 3 of Go/NoGo task, for different valence decoding schemes.

**Figure S9.**
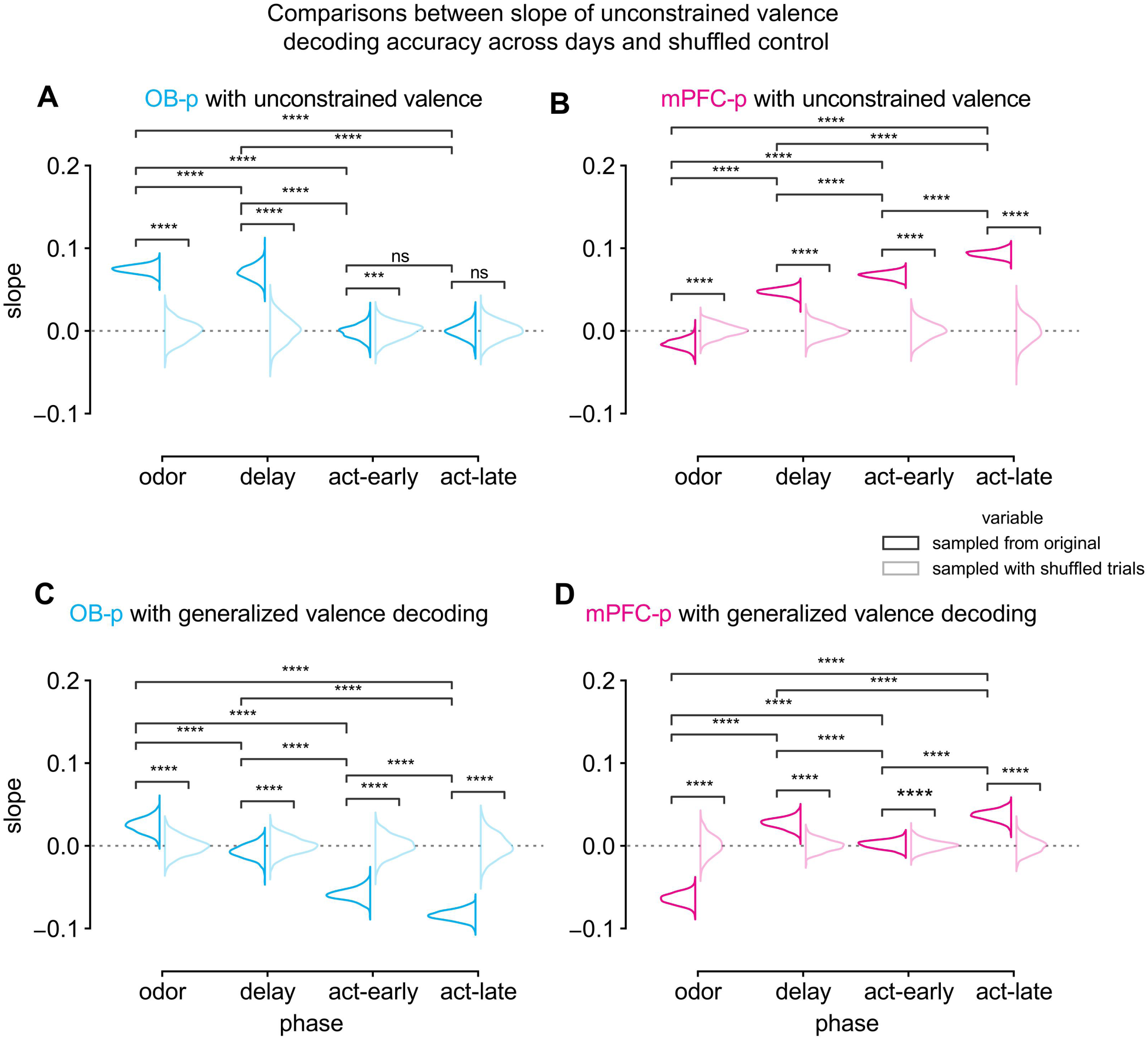
Learning effects on unconstrained and generalized valence decoding in Go/NoGo task. Same as **Figure S7A** but for unconstrained (**Figure S9D**) and generalized (Figure 5D) valence decoding. Due to the imbalanced number of correct trials across animals to construct pseudo-populations, we did not perform similar slope analyses for valence in correct trials. Two-sided Mann-Whitney U tests with Bonferroni corrections were performed.

**Figure S10.**
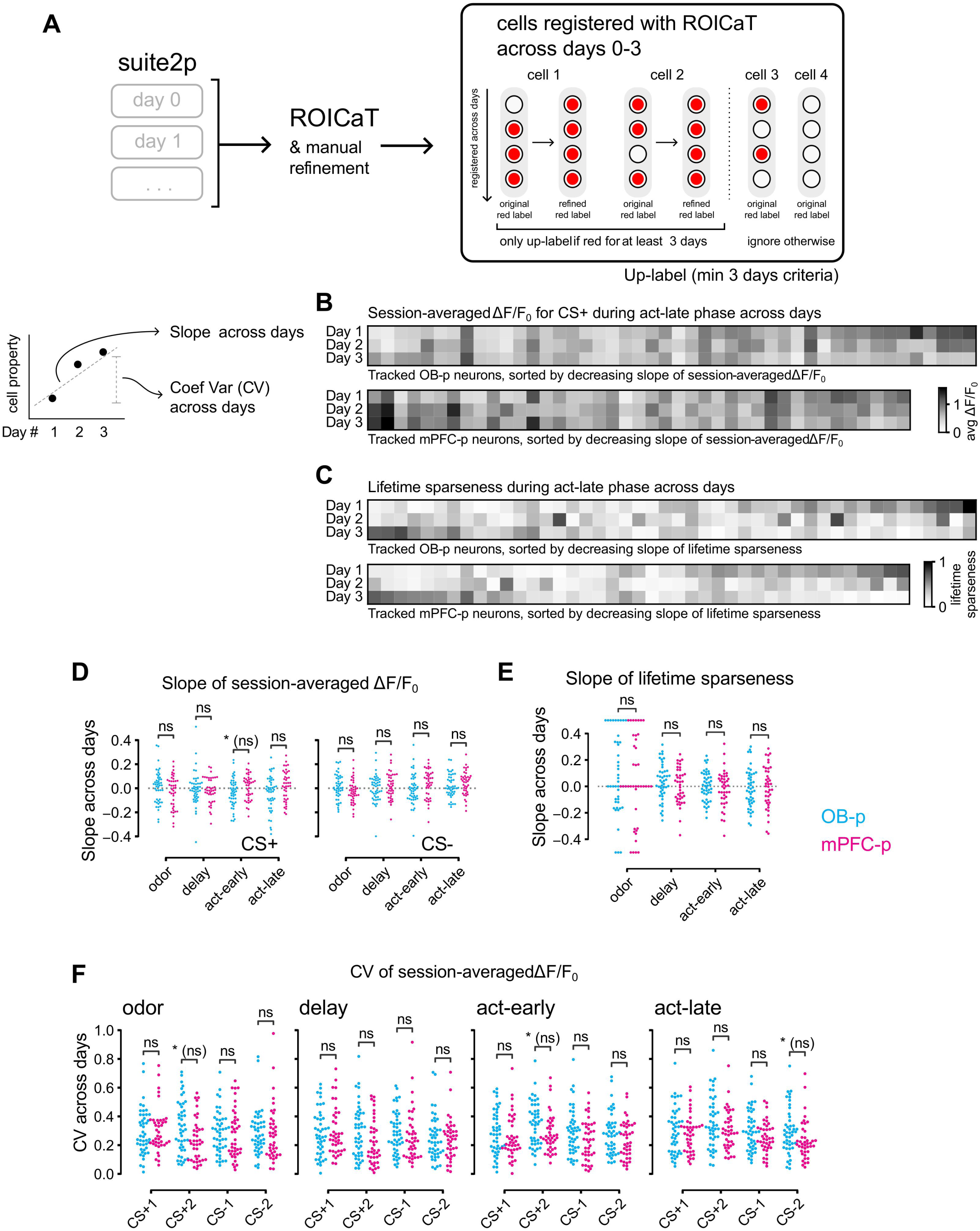
Longitudinal tracking method and session-to-session analyses of cellular properties of projection neurons in Go/NoGo task. (**A**) Schematics of longitudinal tracking using ROICaT after Suite2p segmentation, and up-labeling procedure for projection targets of cells present across all 4 sessions (Days 0-3). Only projection neurons during GNG days were further analyzed in the main text. Tracked neurons that were tagged for at least 3 sessions would be up-labeled, and considered projection neurons for the downstream analyses of tracked projection neurons. (**D, E**) Statistical comparisons of slope across days for single-cell session-averaged normalized fluorescence (**D**) and lifetime sparseness (**E**), between tracked projection neurons (colors) across task phases. (**F**) Comparisons of CV across days for single-cell session-averaged normalized fluorescence, across different odors and phases. All of the comparisons in for panels **D-F** were performed with two-sided Mann-Whitney U tests with Benjamini-Hochberg corrections (ns: p > 0.05; *: 0.01 < p <= 0.05; **: 1 × 10⁻^3^ < p <= 1 × 10⁻^2^; ***: 1 × 10⁻^4^ < p <= 1 × 10⁻^3^; ****: p <= 1 × 10⁻^4^; * (ns): significant with level * before correction but non-significant after correction).

**Figure S11.**
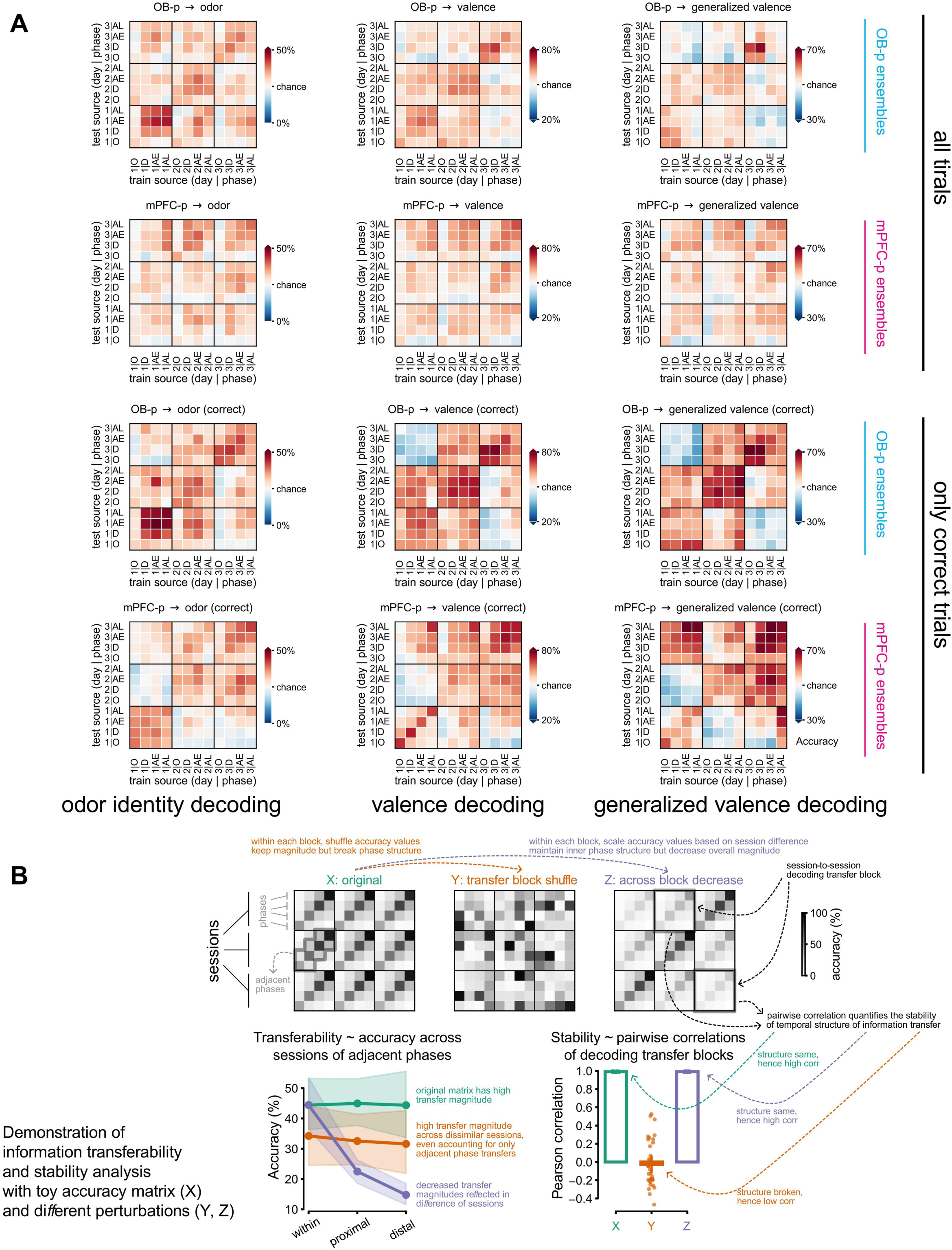
Session transferability matrices across different decoding tasks using tracked projection ensembles, with demonstration of information transferability and stability using toy data. (A) Analysis was performed for each ensemble in each animal separately, and the averaged accuracies across animals are shown in this figure. (**B**) Quantification of transferability and stability of tracked ensemble decoding using toy data. The original toy matrix (**X**, green) is an example of a cross-session and cross-phase decoding accuracy matrix. A decoding transfer block (or “transfer block” for short) is a 4-by-4 matrix that describes how a decoder trained on one set of phases performs when tested on another set, either within or across sessions. Perturbation by shuffling inside each of these blocks (**Y**, orange) would still maintain high cross-session decoding transferability, i.e. the magnitudes of accuracy, but would disrupt the temporal phase-to-phase structure to result in lower information stability. On the other hand, decreasing the overall magnitude for more dissimilar sessions (**Z**, purple) would effectively decrease distal-session transferability, but would still maintain high information stability.

**Figure S12.**
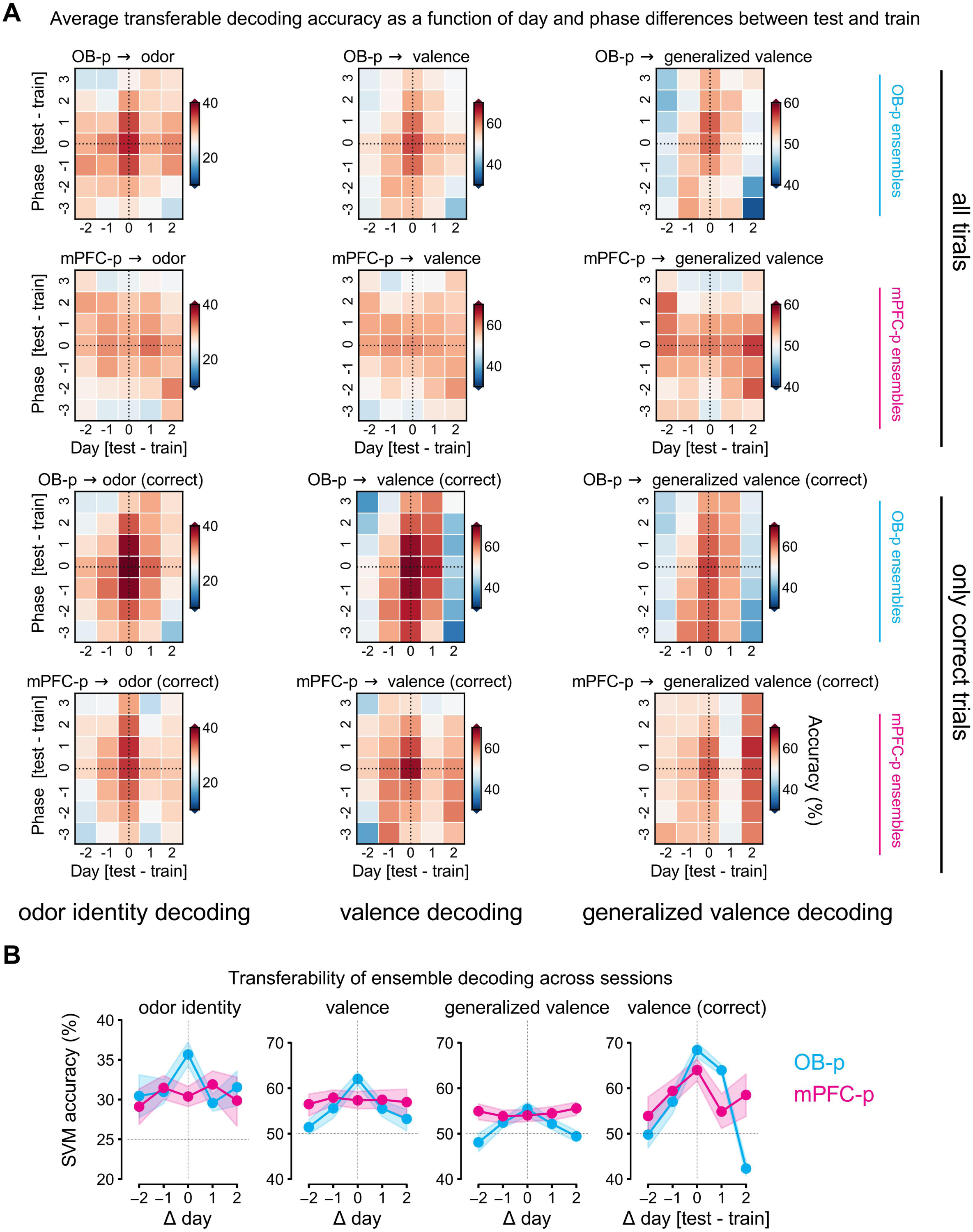
Transferability of ensemble SVM across sessions. (A) Averaged accuracy of SVM trained on different SVM decoding tasks, as a function of day difference (test - train; x-axis) and phase difference (test - train; y-axis). Here, day difference is from numeric values e.g., training on Day 1 then testing Day 3 means difference = 2. Phase difference is from the order of the task phase e.g., training on *act-late* then testing on *odor* means difference = −3. (**B**) Similar to Figure 5F, but showing the directions of train/test day difference. Two-sided Mann-Whitney U tests with Bonferroni corrections were performed.

**Figure S13.**
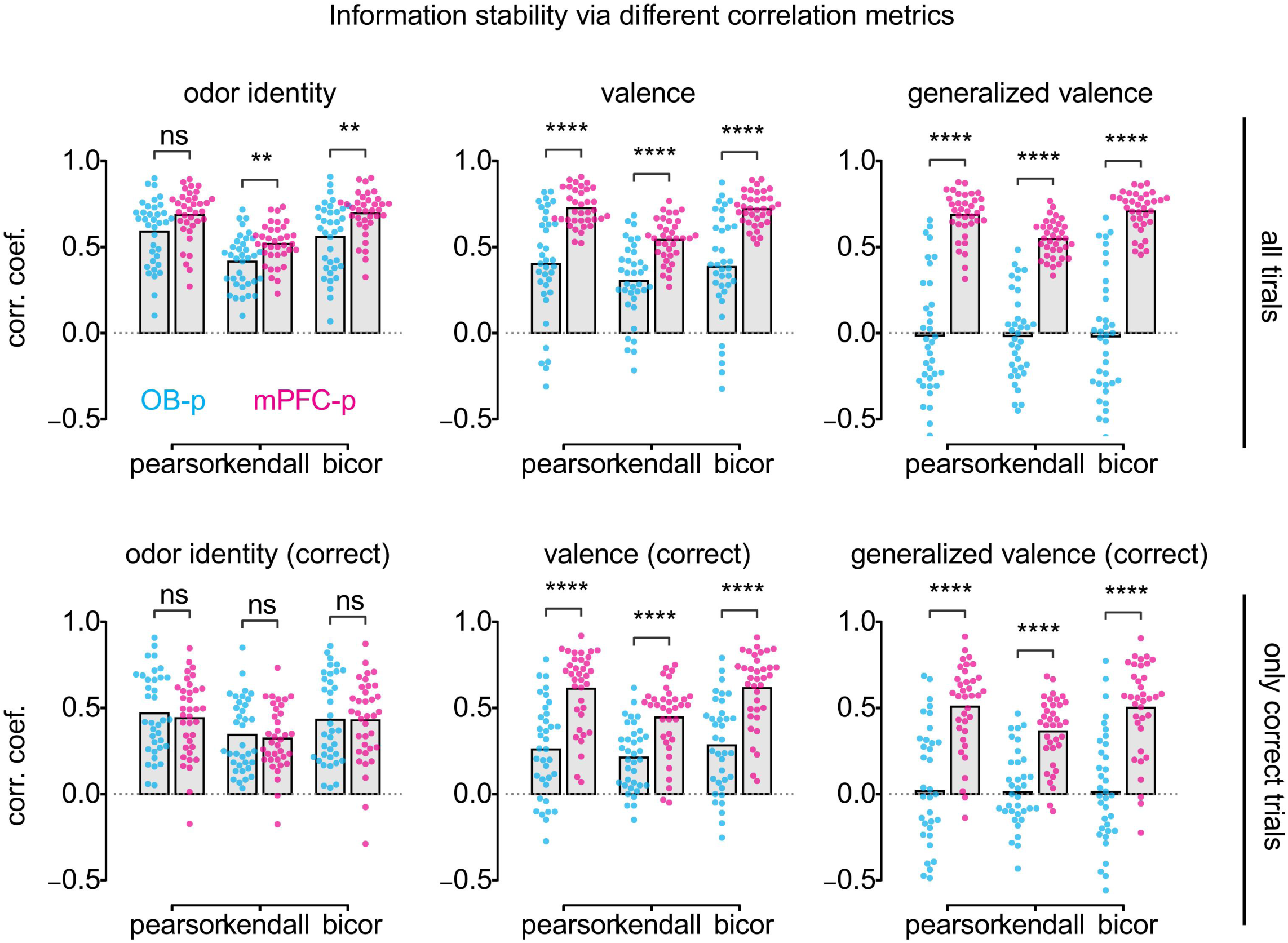
Information stability analyses of tracked ensembles via different correlation methods. Similar to Figure 5G, with all 6 decoding tasks and different correlation methods. Two-sided Mann-Whitney U tests with Bonferroni corrections were performed.

## Notes

### Competing Interest Statement

The authors have declared no competing interest.

## References

1. Hubel DH, Wiesel TN. Receptive fields of single neurones in the cat’s striate cortex. J Physiol. 1959;148(3):574–91.

2. Mountcastle VB. MODALITY AND TOPOGRAPHIC PROPERTIES OF SINGLE NEURONS OF CAT’S SOMATIC SENSORY CORTEX. J Neurophysiol. 1957;20(4):408–34.

3. Felleman DJ, Essen DCV. Distributed Hierarchical Processing in the Primate Cerebral Cortex. Cereb Cortex. 1991;1(1):1–47.

4. Nassi JJ, Callaway EM. Parallel processing strategies of the primate visual system. Nat Rev Neurosci. 2009;10(5):360–72.

5. Hochstein S, Ahissar M. View from the Top Hierarchies and Reverse Hierarchies in the Visual System. Neuron. 2002;36(5):791–804.

6. Iwamura Y. Hierarchical somatosensory processing. Curr Opin Neurobiol. 1998;8(4):522–8.

7. Schreiner CE, Winer JA. Auditory Cortex Mapmaking: Principles, Projections, and Plasticity. Neuron. 2007;56(2):356–65.

8. Spierer L, Lucia MD, Bernasconi F, Grivel J, Bourquin NMP, Clarke S, et al. Learning-induced plasticity in human audition: Objects, time, and space. Hear Res. 2011;271(1–2):88–102.

9. Markov NT, Kennedy H. The importance of being hierarchical. Curr Opin Neurobiol. 2013;23(2):187–94.

10. Kwon SE, Yang H, Minamisawa G, O’Connor DH. Sensory and decision-related activity propagate in a cortical feedback loop during touch perception. Nat Neurosci. 2016;19(9):1243–9.

11. Briggs F. Role of Feedback Connections in Central Visual Processing. Annu Rev Vis Sci. 2020;6(1):1–22.

12. Kreiman G, Serre T. Beyond the feedforward sweep: feedback computations in the visual cortex. Ann N Y Acad Sci. 2020;1464(1):222–41.

13. Asilador A, Llano DA. Top-Down Inference in the Auditory System: Potential Roles for Corticofugal Projections. Front Neural Circuits. 2021;14:615259.

14. Petro LS, Paton AT, Muckli L. Contextual modulation of primary visual cortex by auditory signals. Philos Trans R Soc B Biol Sci. 2017;372(1714):20160104.

15. Elder JH. Shape from Contour: Computation and Representation. Annu Rev Vis Sci. 2018;4(1):423–50.

16. Freiwald WA. The neural mechanisms of face processing: cells, areas, networks, and models. Curr Opin Neurobiol. 2020;60:184–91.

17. Kar K, DiCarlo JJ. Fast Recurrent Processing via Ventrolateral Prefrontal Cortex Is Needed by the Primate Ventral Stream for Robust Core Visual Object Recognition. Neuron. 2021;109(1):164–176.e5.

18. Baxter MG, Gaffan D, Kyriazis DA, Mitchell AS. Orbital Prefrontal Cortex Is Required for Object-in-Place Scene Memory But Not Performance of a Strategy Implementation Task. J Neurosci. 2007;27(42):11327–33.

19. Rao RPN, Ballard DH. Predictive coding in the visual cortex: a functional interpretation of some extra-classical receptive-field effects. Nat Neurosci. 1999;2(1):79–87.

20. Friston K. A theory of cortical responses. Philos Trans R Soc B Biol Sci. 2005;360(1456):815–36.

21. Mori K, Sakano H. How Is the Olfactory Map Formed and Interpreted in the Mammalian Brain? Neuroscience. 2011;34(1):467–99.

22. Wilson DA, Sullivan RM. Cortical Processing of Odor Objects. Neuron. 2011;72(4):506–19.

23. Giessel AJ, Datta SR. Olfactory maps, circuits and computations. Curr Opin Neurobiol. 2013;24(1):120–32.

24. Menelaou G, Diez I, Zelano C, Zhou G, Persson J, Sepulcre J, et al. Stepwise pathways from the olfactory cortex to central hub regions in the human brain. Hum Brain Mapp. 2024;45(18):e26760.

25. Price JL, Powell TP. An experimental study of the origin and the course of the centrifugal fibres to the olfactory bulb in the rat. J Anat. 1970;107(Pt 2):215–37.

26. Haberly LB. Parallel-distributed Processing in Olfactory Cortex: New Insights from Morphological and Physiological Analysis of Neuronal Circuitry. Chem Senses. 2001;26(5):551–76.

27. Chen CFF, Zou DJ, Altomare CG, Xu L, Greer CA, Firestein SJ. Nonsensory target-dependent organization of piriform cortex. Proc Natl Acad Sci. 2014;111(47):16931–6.

28. Diodato A, Brimont MR de, Yim YS, Derian N, Perrin S, Pouch J, et al. Molecular signatures of neural connectivity in the olfactory cortex. Nat Commun. 2016;7(1):12238.

29. Mazo C, Grimaud J, Shima Y, Murthy VN, Lau CG. Distinct projection patterns of different classes of layer 2 principal neurons in the olfactory cortex. Sci Rep. 2017;7(1):8282.

30. Padmanabhan K, Osakada F, Tarabrina A, Kizer E, Callaway EM, Gage FH, et al. Centrifugal Inputs to the Main Olfactory Bulb Revealed Through Whole Brain Circuit-Mapping. Front Neuroanat. 2019;12:115.

31. Chen Y, Chen X, Baserdem B, Zhan H, Li Y, Davis MB, et al. High-throughput sequencing of single neuron projections reveals spatial organization in the olfactory cortex. Cell. 2022;185(22):4117–4134.e28.

32. Chae H, Banerjee A, Dussauze M, Albeanu DF. Long-range functional loops in the mouse olfactory system and their roles in computing odor identity. Neuron. 2022;110(23):3970–3985.e7.

33. Chapuis J, Wilson DA. Bidirectional plasticity of cortical pattern recognition and behavioral sensory acuity. Nat Neurosci. 2012;15(1):155.

34. Kay LM. Circuit oscillations in odor perception and memory. Prog Brain Res. 2014;208:223–51.

35. Jammal L, Whalley B, Ghosh S, Lamrecht R, Barkai E. Physiological expression of olfactory discrimination rule learning balances whole-population modulation and circuit stability in the piriform cortex network. Physiol Rep. 2016;4(14):e12830.

36. Meissner-Bernard C, Dembitskaya Y, Venance L, Fleischmann A. Encoding of Odor Fear Memories in the Mouse Olfactory Cortex. Curr Biol. 2019;29(3).

37. Meissner-Bernard C, Zenke F, Friedrich RW. Geometry and dynamics of representations in a precisely balanced memory network related to olfactory cortex. eLife. 2025;13:RP96303.

38. Dana H, Sun Y, Mohar B, Hulse BK, Kerlin AM, Hasseman JP, et al. High-performance calcium sensors for imaging activity in neuronal populations and microcompartments. Nat Methods. 2019;16(7):649–57.

39. Tervo DGR, Hwang BY, Viswanathan S, Gaj T, Lavzin M, Ritola KD, et al. A Designer AAV Variant Permits Efficient Retrograde Access to Projection Neurons. Neuron. 2016 Oct;92(2):372–82.

40. Madisen L, Zwingman TA, Sunkin SM, Oh SW, Zariwala HA, Gu H, et al. A robust and high-throughput Cre reporting and characterization system for the whole mouse brain. Nat Neurosci. 2010;13(1):133–40.

41. Sattin A, Nardin C, Daste S, Moroni M, Reddy I, Liberale C, et al. Aberration correction in long GRIN lens-based microendoscopes for extended field-of-view two-photon imaging in deep brain regions. eLife. 2025;13:RP101420.

42. Abnar S, Zuidema W. Quantifying Attention Flow in Transformers [Internet]. arXiv; 2020 [cited 2025 Oct 27]. Available from: https://arxiv.org/abs/2005.00928

43. Stettler DD, Axel R. Representations of Odor in the Piriform Cortex. Neuron. 2009;63(6):854–64.

44. Miura K, Mainen ZF, Uchida N. Odor Representations in Olfactory Cortex: Distributed Rate Coding and Decorrelated Population Activity. Neuron. 2012;74(6):1087–98.

45. Roland B, Deneux T, Franks KM, Bathellier B, Fleischmann A. Odor identity coding by distributed ensembles of neurons in the mouse olfactory cortex. eLife. 2017;6:e26337.

46. Bolding KA, Franks KM. Complementary codes for odor identity and intensity in olfactory cortex. eLife. 2017;6:e22630.

47. Nagappan S, Franks KM. Parallel processing by distinct classes of principal neurons in the olfactory cortex. eLife. 2021;10:e73668.

48. Cleland TA, Sethupathy P. Non-topographical contrast enhancement in the olfactory bulb. BMC Neurosci. 2006 Dec;7(1):7.

49. Banerjee A, Marbach F, Anselmi F, Koh MS, Davis MB, Garcia da Silva P, et al. An Interglomerular Circuit Gates Glomerular Output and Implements Gain Control in the Mouse Olfactory Bulb. Neuron. 2015 July;87(1):193–207.

50. Roland B, Jordan R, Sosulski DL, Diodato A, Fukunaga I, Wickersham I, et al. Massive normalization of olfactory bulb output in mice with a “monoclonal nose.” eLife. 2016;5:e16335.

51. Sul JH, Kim H, Huh N, Lee D, Jung MW. Distinct Roles of Rodent Orbitofrontal and Medial Prefrontal Cortex in Decision Making. Neuron. 2010 May;66(3):449–60.

52. Otis JM, Namboodiri VMK, Matan AM, Voets ES, Mohorn EP, Kosyk O, et al. Prefrontal cortex output circuits guide reward seeking through divergent cue encoding. Nature. 2017 Mar;543(7643):103–7.

53. Wang PY, Boboila C, Chin M, Higashi-Howard A, Shamash P, Wu Z, et al. Transient and Persistent Representations of Odor Value in Prefrontal Cortex. Neuron. 2020;108(1):209–224.e6.

54. Zandt EE in ‘t, Cansler HL, Denson HB, Wesson DW. Centrifugal Innervation of the Olfactory Bulb: A Reappraisal. eNeuro. 2019;6(1):ENEURO.0390–18.2019.

55. Zavitz D, Youngstrom IA, Borisyuk A, Wachowiak M. Effect of Interglomerular Inhibitory Networks on Olfactory Bulb Odor Representations. J Neurosci. 2020;40(31):5954–69.

56. Zak JD, Reddy G, Konanur V, Murthy VN. Distinct information conveyed to the olfactory bulb by feedforward input from the nose and feedback from the cortex. Nat Commun. 2024;15(1):3268.

57. Nagayama S, Enerva A, Fletcher ML, Masurkar AV, Igarashi KM, Mori K, et al. Differential Axonal Projection of Mitral and Tufted Cells in the Mouse Main Olfactory System. Front Neural Circuits. 2010;4:120.

58. Franks KM, Russo MJ, Sosulski DL, Mulligan AA, Siegelbaum SA, Axel R. Recurrent Circuitry Dynamically Shapes the Activation of Piriform Cortex. Neuron. 2011;72(1):49–56.

59. Poo C, Isaacson JS. A Major Role for Intracortical Circuits in the Strength and Tuning of Odor-Evoked Excitation in Olfactory Cortex. Neuron. 2011;72(1):41–8.

60. Large AM, Vogler NW, Mielo S, Oswald AMM. Balanced feedforward inhibition and dominant recurrent inhibition in olfactory cortex. Proc Natl Acad Sci. 2016;113(8):2276–81.

61. Bolding KA, Franks KM. Recurrent cortical circuits implement concentration-invariant odor coding. Science [Internet]. 2018;361(6407). Available from: https://app.readcube.com/library/e862fd87-1972-4caf-8c49-302baa8c9bb2/item/3bf0c0e2-42a9-4b53-868d-184b575d1d73

62. Zeppilli S, Ackels T, Attey R, Klimpert N, Ritola KD, Boeing S, et al. Molecular characterization of projection neuron subtypes in the mouse olfactory bulb. eLife. 2021;10:e65445.

63. Makino H, Komiyama T. Learning enhances the relative impact of top-down processing in the visual cortex. Nat Neurosci. 2015;18(8):1116–22.

64. Chen JL, Voigt FF, Javadzadeh M, Krueppel R, Helmchen F. Long-range population dynamics of anatomically defined neocortical networks. eLife. 2016;5:e14679.

65. Allen WE, Kauvar IV, Chen MZ, Richman EB, Yang SJ, Chan K, et al. Global Representations of Goal-Directed Behavior in Distinct Cell Types of Mouse Neocortex. Neuron. 2017;94(4):891–907.e6.

66. Sych Y, Fomins A, Novelli L, Helmchen F. Dynamic reorganization of the cortico-basal ganglia-thalamo-cortical network during task learning. Cell Rep. 2022;40(12):111394.

67. Wilson DA. Habituation of Odor Responses in the Rat Anterior Piriform Cortex. J Neurophysiol. 1998;79(3):1425–40.

68. Ramirez-Gordillo D, Ma M, Restrepo D. Precision of Classification of Odorant Value by the Power of Olfactory Bulb Oscillations Is Altered by Optogenetic Silencing of Local Adrenergic Innervation. Front Cell Neurosci. 2018;12:48.

69. Wang D, Liu P, Mao X, Zhou Z, Cao T, Xu J, et al. Task-Demand-Dependent Neural Representation of Odor Information in the Olfactory Bulb and Posterior Piriform Cortex. J Neurosci. 2019;39(50):10002–18.

70. Kudryavitskaya E, Marom E, Shani-Narkiss H, Pash D, Mizrahi A. Flexible categorization in the mouse olfactory bulb. Curr Biol. 2021;31(8):1616–1631.e4.

71. Hernandez DE, Ciuparu A, Silva PG da, Velasquez CM, Rebouillat B, Gross MD, et al. Fast updating feedback from piriform cortex to the olfactory bulb relays multimodal identity and reward contingency signals during rule-reversal. Nat Commun. 2025;16(1):937.

72. Liu D, Gu X, Zhu J, Zhang X, Han Z, Yan W, et al. Medial prefrontal activity during delay period contributes to learning of a working memory task. Science. 2014;346(6208):458–63.

73. Falco ED, An L, Sun N, Roebuck AJ, Greba Q, Lapish CC, et al. The Rat Medial Prefrontal Cortex Exhibits Flexible Neural Activity States during the Performance of an Odor Span Task. eNeuro. 2019;6(2):ENEURO.0424–18.2019.

74. Jun H, Lee JY, Bleza NR, Ichii A, Donohue JD, Igarashi KM. Prefrontal and lateral entorhinal neurons co-dependently learn item–outcome rules. Nature. 2024;1–8.

75. Yang J, Quraish AU, Murakami K, Ishikawa Y, Takayanagi M, Kakuta S, et al. Quantitative analysis of axon collaterals of single neurons in layer IIa of the piriform cortex of the guinea pig. J Comp Neurol. 2004;473(1):30–42.

76. Chéreau R, Williams LE, Bawa T, Holtmaat A. Circuit mechanisms for cortical plasticity and learning. Semin Cell Dev Biol. 2022;125:68–75.

77. Feulner B, Perich MG, Miller LE, Clopath C, Gallego JA. A neural implementation model of feedback-based motor learning. Nat Commun. 2025;16(1):1805.

78. Pachitariu M, Stringer C, Dipoppa M, Schröder S, Rossi LF, Dalgleish H, et al. Suite2p: beyond 10,000 neurons with standard two-photon microscopy [Internet]. Neuroscience; 2016 [cited 2025 Dec 5]. Available from: http://biorxiv.org/lookup/doi/10.1101/061507

79. Friedrich J, Zhou P, Paninski L. Fast Online Deconvolution of Calcium Imaging Data. arXiv. 2016;13(3):e1005423.

80. Dosovitskiy A, Beyer L, Kolesnikov A, Weissenborn D, Zhai X, Unterthiner T, et al. An Image is Worth 16×16 Words: Transformers for Image Recognition at Scale [Internet]. arXiv; 2020 [cited 2025 Oct 27]. Available from: https://arxiv.org/abs/2010.11929

81. Kingma DP, Ba J. Adam: A Method for Stochastic Optimization [Internet]. arXiv; 2014 [cited 2025 Oct 27]. Available from: https://arxiv.org/abs/1412.6980

82. Srinivasan S, Daste S, Modi MN, Turner GC, Fleischmann A, Navlakha S. Effects of stochastic coding on olfactory discrimination in flies and mice. PLOS Biol. 2023;21(10):e3002206.

